# Dopamine Signaling in the Dorsomedial Striatum Promotes Compulsive Behavior

**DOI:** 10.1101/2020.03.30.016238

**Authors:** Jillian L. Seiler, Caitlin V. Cosme, Venus N. Sherathiya, Joseph M. Bianco, Abigael S. Bridgemohan, Talia N. Lerner

## Abstract

Compulsive behavior is a defining feature of disorders such as substance use disorder and obsessive-compulsive disorder. Current evidence suggests that corticostriatal circuits control the expression of established compulsions, but little is known about the mechanisms regulating the development of compulsions. We hypothesized that dopamine, a critical modulator of striatal synaptic plasticity, could control alterations in corticostriatal circuits leading to the development of compulsions (defined as continued reward-seeking in the face of punishment). We used dual-site fiber photometry to measure dopamine axon activity in the dorsomedial striatum (DMS) and the dorsolateral striatum (DLS) as compulsions emerged. Individual variability in the speed with which compulsions emerged was predicted by DMS dopamine axon activity. Amplifying this dopamine signal accelerated animals’ transitions to compulsion, whereas inhibition led to learning delays. In contrast, amplifying DLS dopamine signaling had no effect on the emergence of compulsions. These results establish DMS dopamine signaling as a key controller of the development of compulsive reward-seeking.

## INTRODUCTION

Animals learn about the consequences of their actions through reinforcement. Positive or negative outcomes lead to the formation of action-outcome associations, which allow an animal to predict consequences and act purposefully. Action-outcome learning relies on the dorsomedial striatum (DMS) and supports goal-directed behavior (Yin, Knowlton, et al., 2005; Yin, Ostlund, et al., 2005). Goal-directed behavior has advantages and disadvantages. Its chief benefit is that it permits behavioral flexibility; a goal-directed animal can readily adapt to new circumstances. However, flexibility comes with two costs. First, goal-directed behavior demands cognitive resources to execute actions. Second, excessive behavioral flexibility might lead to animals prematurely abandoning strategies that would be productive in the longer run, as when action-outcome associations fluctuate or are probabilistic. Such situations often occur in nature.

Given these disadvantages, animals have developed brain systems that support the continuation of ingrained behavioral patterns even when action-outcome contingencies change. Habits, which rely on the dorsolateral striatum (DLS; Yin et al., 2004), decouple actions from outcomes and instead promote the control of behavior through stimulus-response associations (Yin & Knowlton, 2006). The transition to habitual responding during prolonged training can be measured as an insensitivity to outcome devaluation or as an insensitivity to changes in action-outcome contingency (Rossi & Yin, 2012; Yin & Knowlton, 2006). Compulsive behavior may or may not be an extreme form of habit, in which reward-seeking behaviors continue in the face of punishment (Gillan et al., 2016; Lipton et al., 2019; Lüscher et al., 2020). Compulsive behavior, herein referred to specifically as “punishment-resistant reward-seeking,” like habit, appears to depend on dorsal striatal brain regions and their cortical inputs (Lipton et al., 2019; Lüscher et al., 2020; Lüscher & Janak, 2021), but little is known about how compulsions emerge (Lüscher & Janak, 2021).

One hypothesis regarding the relationship between habits and punishment-resistant reward-seeking is that a reliance on habit makes it generally difficult for an animal to adjust its reward-seeking behavior when action-outcome associations change, including in circumstances in which previously rewarded actions begin to produce negative outcomes (Giuliano et al., 2019; Lüscher et al., 2020). In this case, the development of punishment-resistant reward-seeking relies on the DLS and habit formation precedes the emergence of this behavior. In accord with this idea, the emergence of DLS dopamine signaling and the dependence of reward-seeking behavior on DLS dopamine seem to precede the development of punishment-resistant reward-seeking for addictive drugs (Giuliano et al., 2019; Willuhn et al., 2012). However, habit formation is not *required* for punishment-resistant reward-seeking to emerge. For example, when Singer et al (2017) trained rats to perform new action sequences each day to get cocaine, punishment-resistant drug-seeking developed that was independent of habit and DLS dopamine signaling.

An alternative hypothesis is that punishment-resistant reward-seeking arises due to strengthened action-outcome circuits centered on the DMS. Enhanced activity in orbitofrontal cortex (OFC) to DMS projections has been observed in animals that compulsively self-stimulate their VTA dopamine neurons and in animals that self-administer methamphetamine in a punishment-resistant way (Hu et al., 2019a; Pascoli et al., 2015, 2018). However, it is unclear what mechanisms might lead to a strengthening of OFC-DMS connections.

In this study, we trained mice on a random interval schedule (RI60) that promotes habit formation and tested whether it also led to punishment-resistant reward-seeking. We found that prolonged RI60 training did lead to punishment-resistant reward-seeking, although, interestingly, only in a subset of mice. RI60 training also led to an increase in the time it took mice to learn a reversed action-outcome contingency (“omission”), which is a measure of habitual, inflexible responding that does not depend on punishment. The fact that RI60 training increased both types of inflexible responding on a group level, but not always in the same animals, led us to hypothesize that the neural circuits that drive the development of punishment-resistant reward-seeking may be distinct from those driving habit formation.

Differences in dorsal striatal dopamine signaling could contribute to individual differences in reward learning trajectories related to the development of punishment-resistance as opposed to simple habit. Dopamine is well-known to regulate reward learning through its encoding of reward prediction error, and it operates as a critical neuromodulator in both the DMS and DLS (Kreitzer & Malenka, 2008; Lerner et al., 2021; Lovinger, 2010; Nicola et al., 2000; Schultz et al., 1997). Dopamine receptor blockade in the DMS can inhibit action-outcome learning (Yin et al., 2009), while dopamine signaling in the DLS is required for habit formation (Faure et al., 2005). Blocking dopamine receptors in the DLS can also inhibit drug-seeking for cocaine or alcohol (Corbit et al., 2012; Hodebourg et al., 2018; Murray et al., 2012; Pacchioni et al., 2011; Vanderschuren et al., 2005).

To examine changes in dopamine activity in the DMS and DLS over the course of learning, we used dual-site fiber photometry to record calcium transients in dopamine axons in both of these regions. We found that the DMS but not the DLS dopamine axon signal predicts whether an individual will continue to pursue sucrose rewards even when faced with the possibility of a footshock punishment. We then optogenetically manipulated the DMS dopamine signal and confirmed the causal and temporally-specific role of DMS dopamine in action-outcome learning and the development of punishment-resistant reward-seeking. The optogenetic acceleration of punishment-resistant reward-seeking was not systematically accompanied by an increased resistance to adapting to an action-outcome contingency change in an omission probe, where no positive punishments were delivered. These data suggest that the formation of habits and the development of compulsions rely on distinct dopaminergic reinforcement signals, with the emergence of compulsion driven primarily by DMS dopamine activity.

## RESULTS

### A random interval, but not random ratio, schedule of reinforcement promotes punishment-resistant reward-seeking

We first determined whether training paradigms used to elicit habitual responding also elicit punishment-resistant reward-seeking. Previous literature has demonstrated that a random interval (RI60) schedule of reinforcement, but not a random ratio (RR20) schedule, promotes habit (Derusso et al., 2010; Gremel & Costa, 2013; Wiltgen et al., 2012; Yin, Ostlund, et al., 2005). We therefore compared training on RI60 and RR20 reinforcement schedules to assess whether these schedules have differential effects on punishment-resistant reward-seeking. After initial magazine training and pre-training on a fixed ratio (FR1) schedule, mice were transitioned to either RI30 or RR10 schedules, and then finally to RI60 or RR20 schedules (Fig. 1A). We performed probes for punishment-resistant reward-seeking at an early and late time point: the first after 1-2 days of RI60/RR20 training and the second after 13-14 days of RI60/RR20 training. During the shock probe sessions, nosepokes were accompanied by a ⅓ risk of mild shock (0.2mA, 1s; Fig. 1B). The shock intensity was chosen based on previous studies of punishment-resistant reward-seeking (Harada et al., 2019). We verified that this shock intensity is aversive to the mice in a fear conditioning paradigm in which 12 tone-shock pairings were delivered. The next day, mice that had received tone-shock pairings showed increased freezing to the tone compared with mice that had only been exposed to the tone on the previous day (Fig. S1A; unpaired t test, p<0.01). To test whether punishment-resistant reward-seeking developed in tandem with another test of habit-like behavioral inflexibility, a subset of mice were also tested at the end of training on an omission probe (Fig. 1A; Derusso et al., 2010; Rossi & Yin, 2012; Yu et al., 2009). In the omission probe, mice were required to withhold nosepokes to receive rewards, reversing the previously learned contingency (Fig. 1C).

**Figure 1.**
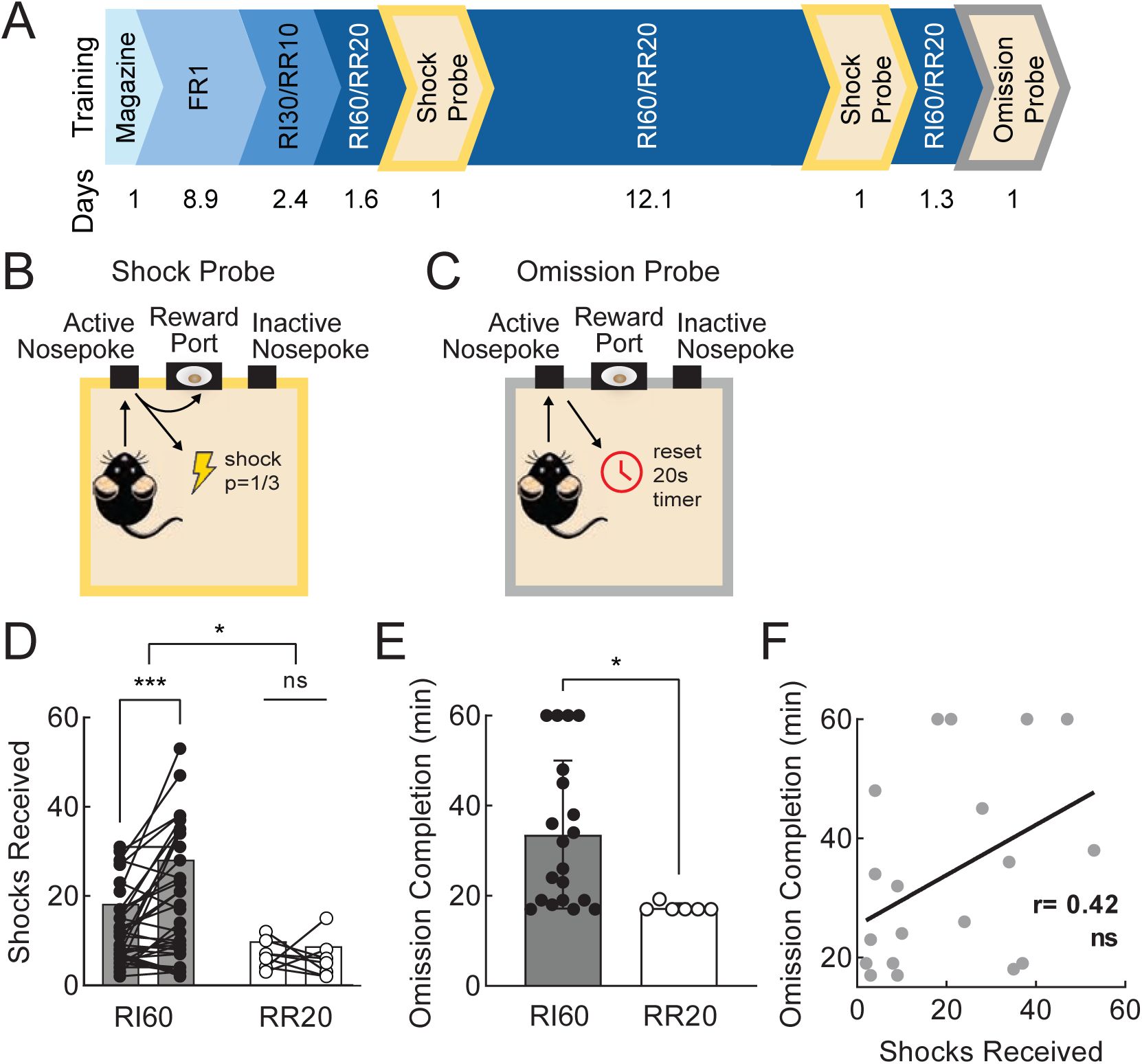
A random interval, but not random ratio, schedule of reinforcement promotes compulsive reward-seeking. **A.** Timeline of operant training and probes. Animals progressed from a fixed ratio to a random ratio or random interval schedule of reinforcement. A shock probe occurred both early and late in RI60/RR20 training and an omission probe occurred at the end of training. Average number of days in each stage of training is given below. **B.** Schematic of shock probe: active nosepokes had a ⅓ probability of incurring a shock consequence. **C.** Schematic of omission probe: rewards were delivered when mice refrained from nosepoking at the previously active port for 20s. **D.** Shocks received on early and late shock probes for RI60-trained (black; n=36) and RR20-trained (white; n=8) mice. Bars represent mean; points represent individuals. *p<0.05, ***p<0.001. **E.** Average time to complete the omission probe (earn 50 rewards; max 60 minutes) for RI60-trained (black; n=20) and RR20-trained (white; n=7) mice. Error bars represent SD. *p<0.05 **F.** Correlation between shocks received on the late shock probe and omission completion time for all RI60-trained mice tested in both probes (r=0.42, ns). See also Supplemental Figure 1.

We observed a significant main effect of schedule (RI60 vs RR20) on the number of shocks mice were willing to receive on the shock probes (two-way ANOVA, F_1,45_=6.31, p<0.05) as well as an interaction of schedule and training time (F_1,45_=4.54, p<0.05). Furthermore, after extended training on the RI60, but not the RR20, schedule, mice increased the number of shocks they were willing to receive (Fig. 1D; Bonferroni, p <0.001). During the second shock probe, both RI60- and RR20-trained mice initially continued to nosepoke at the same rates relative to their training baseline, but the RR20-trained mice reduced their nosepoke rates to near zero by the end of the session (Fig. S1B; two-way ANOVA, main effect of training time, F_4.426,181.5_=4.99, p<0.001, Bonferroni, p<0.05 for bins after 50 minutes). The RR20-trained mice were also more willing than RI60-trained mice to explore alternative actions during the second shock probe session, as indicated by a higher fraction of nosepokes made at the inactive port (Fig. S1D-E; unpaired t-test, p<0.05).

On the omission probe, RI60-trained mice took longer to complete the session than mice trained on RR20 (Fig. 1E; unpaired t-test, p<0.05). RR20-trained mice almost immediately stopped nosepoking during the omission probe session, whereas RI60-trained mice continued to nosepoke for much longer periods of time, albeit with large variability between individuals (Fig. S1C; two-way ANOVA, interaction of schedule and time, F_11,275_=2.22, p<0.05). RR20-trained mice therefore maintain a higher level of flexibility in their behavior when presented with positive punishments and when presented with a reversed contingency as compared to RI60-trained mice.

It is not clear what differences between RI60 and RR20 training might elicit differences in the development of punishment-resistant or omission-resistant reward-seeking. As previously reported (Gremel & Costa, 2013; Wiltgen et al., 2012), RI60 and RR20 training schedules provoked approximately equivalent rates of nosepoking (Fig. S1G). However, we also observed that RI60-trained mice made fewer nosepokes per reward (Fig. S1H; mixed-effects analysis, F_1,42_=21.70, p<0.0001) and earned significantly more rewards per training session, on average, than RR20-trained mice (Fig. S1I; unpaired t-test, p<0.0001). Therefore, these different task structures have different effort demands and incur different reward histories, which could influence learning trajectories.

In both the shock and omission probes, RI60-trained mice showed significant individual variability (omission F-test to compare variances, F_19,5_=305.90, p<0.0001; shock 1 F test to compare variances F_37,8_=5.88, p<0.01; shock 2 F test to compare variances F_37,7_= 12.08, p<0.001). The variability in punishment-resistance was not due to variation in body weight (Fig. S1J). We wondered whether the same individuals who withstood a high number of shocks also took longer to learn the omission contingency. Looking at data from individual animals tested on both probes, we found that these two measures were not significantly correlated (Fig. 1F; r=0.42, ns). Thus, although the RI60 schedule promotes inflexible behavior in the form of both punishment-resistant and omission-resistant reward-seeking, these two phenomena do not necessarily occur in the same individuals and their development may rely on different brain circuits.

### Three behavioral phenotypes emerge with extended RI60 training

The large variation in behavior induced by RI60 training caused us to wonder whether individual mice were taking different strategies to “solve” the RI60 task, which could lead to differences in the development of punishment-resistant reward-seeking. To analyze whether differences in performance in the shock probes were associated with different behavior in the RI60 task, we divided the RI60-trained mice into three analysis groups based on shock probe performance: “punishment resistant” (PR) mice that tolerated a high level of shocks in both the first and second probes (25% of mice), “delayed punishment resistant” (DPR) mice that increased the number of shocks they would tolerate from the first to the second probe (25% of mice), and “punishment sensitive” (PS) mice that would not tolerate many shocks on either the first or second probes (50% of mice; Fig. 2A; see methods for sorting criteria). As expected based on this sorting, a mixed effects analysis showed a significant interaction of phenotype and training time (F_3, 40_ = 24.18, p<0.0001), with a Bonferroni test revealing that only DPR mice showed a significant difference in shocks received between the early and late shock probes (Fig. 2B ; p<0.0001).

**Figure 2.**
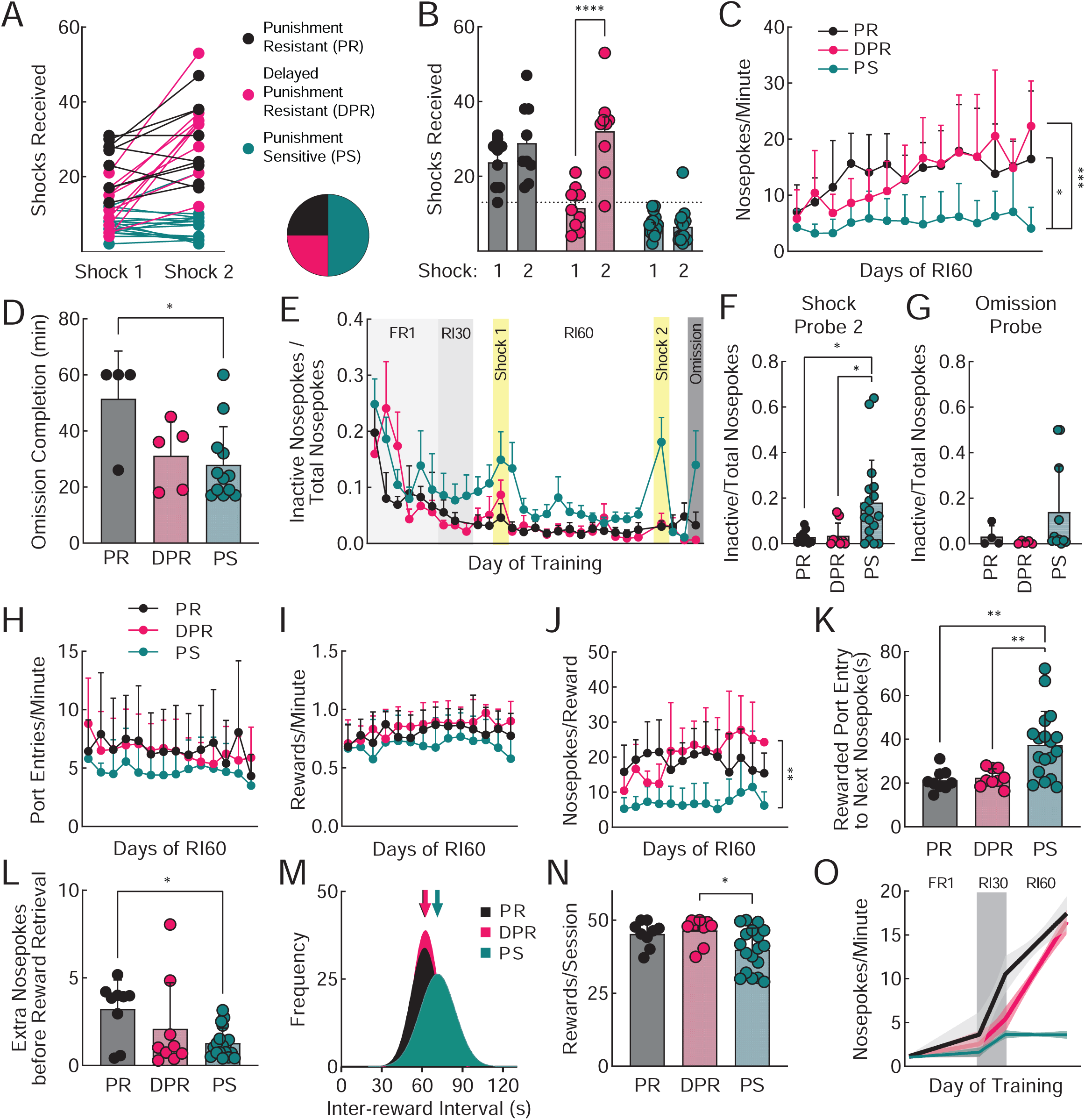
Three punishment-related phenotypes emerge with extended RI60 training. **A.** Shocks received by mice trained under RI60 only (same data as shown in Fig. 1D). Mice were classified as punishment resistant (PR; black), delayed punishment resistant (DPR; pink), or punishment sensitive (PS; teal) based on the number of shocks they received during the early and late shock probes (see Methods). **B.** Average shocks received on early and late shock probes for each phenotype. Error bars represent SD ****p<0.0001. **C.** Average nosepokes per minute across days of RI60 training. Error bars represent SD, PR vs PS: *p<0.01, DPR vs PS: ***p<0.001. **D.** Average time to complete the omission probe (earn 50 rewards; max 60 minutes). Error bars represent SD *p<0.05. **E.** Average nosepokes on the inactive port as a fraction of the total nosepokes across days of training and probe sessions. Error bars represent SD. **F.** Average nosepokes on the inactive port as a fraction of the total nosepokes during the second shock probe session. Error bars represent SD *p<0.05. **G.** Average nosepokes on the inactive port as a fraction of the total nosepokes during the omission probe session. Error bars represent SD. **H.** Average number of port entries per minute across days of RI60 training. Error bars represent SD. **I.** Average number of rewards earned per minute across days of RI60 training. Error bars represent SD. **J.** Average nosepokes made per reward across days of RI60 training. Error bars represent SD **p<0.01. **K.** Average time from a rewarded port entry to the next nosepoke. Error bars represent SD **p<0.01. **L.** Average unrewarded nosepokes made following a rewarded nosepoke, prior to a rewarded port entry. Error bars represent SD *p<0.05. Distribution of inter-reward interval times in seconds for each group. Arrows represent mean. Average rewards earned per RI60 session. Error bars represent SD *p<0.05. **O.** Segmental linear regression showing the slope of nosepokes made per minute in FR1, RI30, and RI60 schedules. Shaded region represents 95% confidence bands. See also Supplemental Figure 2.

When mice were divided into these groups, an analysis of behavior in RI60 sessions showed interesting differences (Figs. 2, S2). Both PR and DPR mice had higher rates of nosepoking than PS mice (Fig. 2C; mixed effects analysis, main effect of training time F_5.236,156.3_=9.79, p<0.0001, main effect of phenotype F_2,33_=18.59, p <0.0001, interaction F_26,388_=3.28, p<0.0001). PR mice also took significantly longer to complete the omission probe in comparison to PS mice (Fig. 2D; one-way ANOVA, F_2,18_=4.36, p<0.05, Tukey’s multiple comparison, PR=51.50 ± 17s vs PS=27.92 ± 13.65s, p<0.05). Meanwhile, PS mice were more likely to explore the inactive nosepoke at the end of training and during the second shock probe session (Fig. 2E; mixed-effects analysis, main effect of phenotype, F_2,33_=3.79, Tukey’s multiple comparisons test, RI60 days 13 and 14 for PS vs DPR, p<0.05; Fig. 2F; one-way ANOVA for shock 2, F_2,18_=5.36, Tukey’s multiple comparison, for PS vs DPR, p<0.05, second shock for PS vs PR, p<0.05; Fig. 2G; no significant effect during omission).

At least some of the variation between the PR, DPR and PS mice was attributable to sex differences. PR mice were more likely to be male, whereas PS mice were more likely to be female. The DPR group was evenly split between the sexes (Fig. S2A). In general, as a group, male mice tolerated more shocks on both shock probes than female mice (Fig. S2B) and had higher nosepoke rates during RI60 (Fig. S2C). However, variance in punishment-resistant reward-seeking was not fully explained by sex. A three-way ANOVA of phenotype x sex x time revealed that more of the variance in shocks received was accounted for by phenotype than by sex (15.09 vs 0.03%, respectively). Nevertheless, given these sex differences, we were careful to include a balance of both male and female mice going forward in all our experiments.

Port entry rates and rates of rewards earned per minute did not differ among the PR, DPR, and PS groups (Fig. 2H-I). As a result, PS mice are more “efficient,” making fewer nosepokes per reward than PR and DPR mice (Fig. 2J; mixed-effects analysis, main effect of training time F_4.65,87.86_=2.42, p<0.05, main effect of phenotype F_2,21_=22.77, p<0.0001, interaction F_24,22_7=1.76, p<0.05). To achieve such efficiency, a mouse needs to understand when a nosepoke is likely to lead to reward. In the RI60 task, a mouse is unlikely to receive a reward if it has recently received one (on average, it will have to wait 60 seconds before its efforts become fruitful again). PS mice waited an average of 38 ± 15 seconds to resume nosepoking after retrieving a reward, significantly longer than the other groups (Fig. 2K; Tukey’s multiple comparison, PR=21 ± 5, p<0.01; DPR=22 ± 4, p<0.01). We also looked at whether mice made “extra” nosepokes after a rewarded nosepoke, before going to the port to collect their reward. PS mice made significantly fewer extra nosepokes before reward retrieval than PR mice (Fig. 2L; one-way ANOVA, F_2,33_=4.24, p<0.05, Tukey’s multiple comparison, PS=1.28 ± 0.85, PR= 3.23 ± 1.65, p<0.05, DPR= 2.1 ± 2.63), which helped maximize the efficiency of their behavior.

Why would PR and DPR mice expend more effort than necessary to earn rewards? On an RI60 reinforcement schedule, the average interval between available rewards is 60 seconds, but there is a normal distribution around this average. To ensure a nosepoke is likely to be rewarded, a mouse should wait to nosepoke, but to “catch” all instances when reward becomes available in the shortest possible amount of time, mice must nosepoke constantly. In other words, if mice are willing to expend the effort, high rates of nosepoking yield rewards at shorter intervals. Indeed, when we looked at the distribution of inter-reward intervals for PR, DPR and PS mice, we found that the inter-reward intervals for PS mice skew longer (Fig. 2M). The average inter-reward interval for PR is 71.2 ± 32.3 seconds and for DPR mice is 69.58 ± 35.18 seconds, while the average for PS mice is 79.77 ± 26.54 seconds (K-S Test, PR vs PS, p<0.0001; DPR vs PS, p<0.0001). PR and DPR mice thus are better able to track the probability of reward availability imposed in the task, whereas PS mice have longer lags before they earn available rewards. Although we did not observe differences in the total number of rewards earned per minute on any particular day of RI60 training (Fig. 2I), when the number of rewards earned per session was averaged over all days, we found that PS received slightly fewer rewards (Fig. 2N; one-way ANOVA, F_2,33_=4.45, p <0.05, Tukey’s multiple comparison, PS= 39.91 ± 7.39, PR= 45.33 ± 4.33, p<0.05). This difference illustrates the advantage of the high-nosepoking approach: maximize rewards at any cost.

We observed that PR and DPR mice take similar reward-seeking strategies in the RI60 task, so to examine differences between them, we looked at earlier training data from FR1 and RI30 sessions. We observed that PR mice escalate their nosepoking more rapidly than DPR mice, as soon as they enter RI30 training (the first RI schedule they encounter; Fig. 2O). DPR mice then escalate their nosepoking throughout RI60 training, concurrent with the development of punishment-resistance. Thus, the timing of nosepoke escalation is a key behavioral predictor of mice who will show punishment-resistant reward-seeking when probed. Importantly, this escalation emerges in PR mice prior to any experience of punishment, suggesting that some mice may have a predisposition towards developing punishment-resistance that is related to their initial reward-seeking strategy.

### Dopamine axon signals in the DMS predict punishment-resistant reward-seeking

To better understand the neural circuits mediating differences in the development of punishment-resistant reward-seeking, we recorded the activity of dopamine axons in the dorsal striatum during RI/RR training. Dopaminergic projections to the DMS and DLS are distinct, meaning that dopamine-mediated reinforcement learning can be separately effectuated in these two areas (Ikemoto, 2007; Lerner et al., 2015). We reasoned that by examining the activity of dopaminergic projections to the DMS and DLS during RI/RR training, we could assess whether different dopamine signals were contributing to different aspects of behavior. To record the activity of dopamine axons in the DMS and DLS in freely moving mice, we injected an adeno-associated virus (AAV) expressing cre-dependent GCaMP7b (AAV5-CAG-FLEX-jGCaMP7b-WPRE) into the substantia nigra pars compacta (SNc) of DAT-IRES-cre mice. We then implanted fiber optic probes above the DMS and DLS to record the activity of dopaminergic axons (Fig. 3A, 4A). Dopaminergic axon activity was recorded simultaneously in the DMS and DLS in all mice. Thus, we could be certain that any differences we observed in activity in the two regions were not due to differences in behavior between groups of mice.

**Figure 3.**
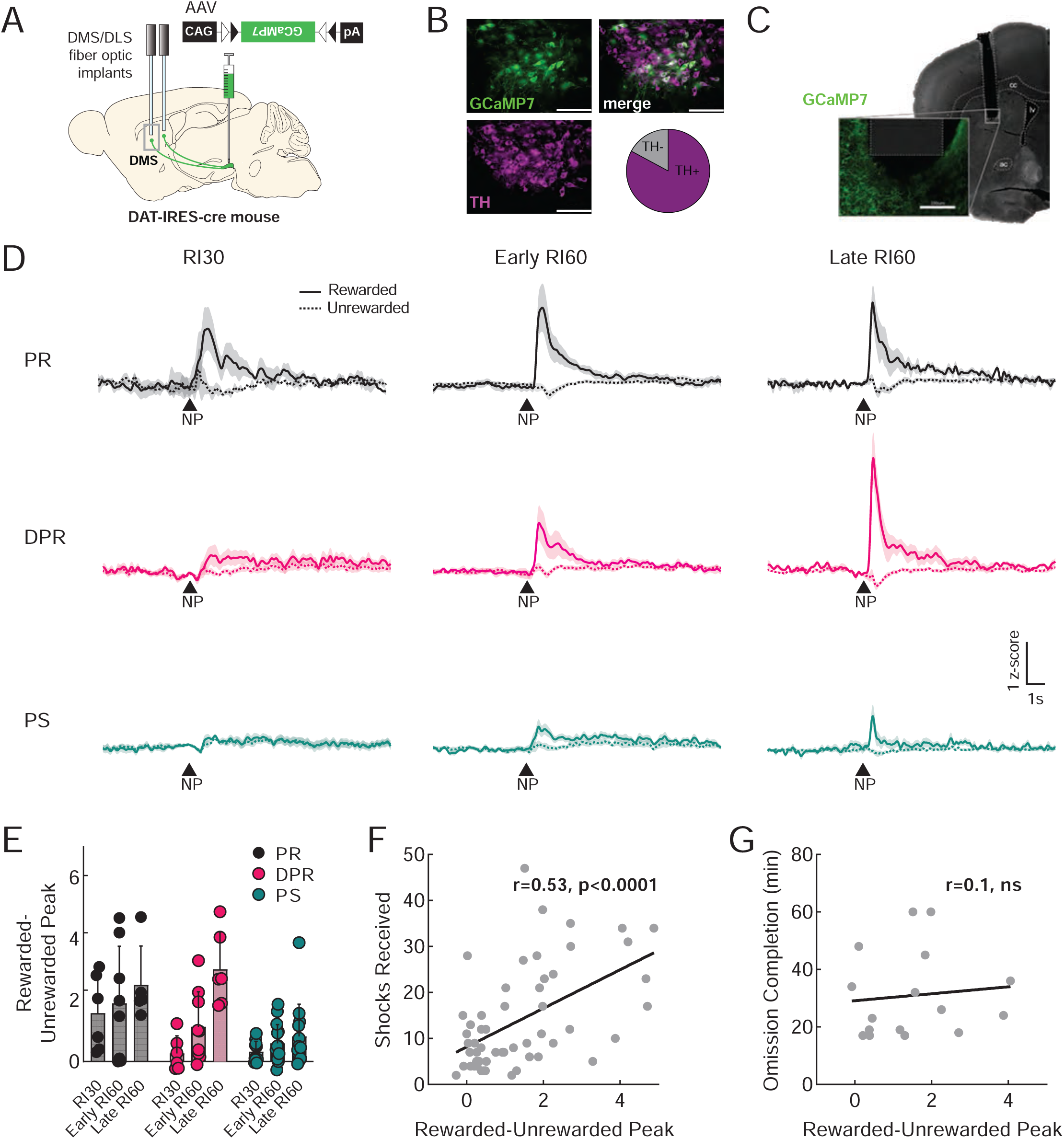
Dopamine axon signals in the DMS predict punishment-resistant reward-seeking. **A.** Viral injection and probe placement strategy for recording from dopamine terminals in the DMS and DLS. DMS is highlighted here. **B.** High magnification (40x) images of SNc showing GCaMP7b expression in green, TH positive cells in magenta, and the merged image. Scale bars are 100μm. Quantification of GCaMP7b-expressing cells that are TH+ is shown; n=572 cells. **C.** Representative image showing probe placement in DMS. Area of magnification shows GCaMP7b expression in dopaminergic axons near the probe site in green. Scale bar is 100μm. cc=corpus callosum, lv=lateral ventricle, ac=anterior commissure. **D.** Peri-stimulus time histograms (PSTHs) showing the average signal from DMS dopamine terminals at the time of rewarded (solid) and unrewarded (dashed) nosepokes (NP) for each phenotype during RI30 training, early in training (RI60 day one or two), and late in training (RI60 day eleven or twelve). Shaded region represents SEM. Punishment resistant (PR; black), delayed punishment resistant (DPR; pink), or punishment sensitive (PS; teal). **E.** Quantification of the average “difference score” for DMS dopamine terminal signals observed in response to rewarded and unrewarded nosepokes (calculated as the peak at the time of rewarded nosepokes minus the peak at the time of unrewarded nosepokes). Error bars represent SD. **F.** Correlation of shocks received in shock probe sessions and “difference score” in DMS dopamine terminals (r=0.53, p<0.0001). **G.** Correlation of time to complete omission and “difference score” in DMS dopamine terminals (r=0.1, ns). See also Supplemental Figure 3.

**Figure 4.**
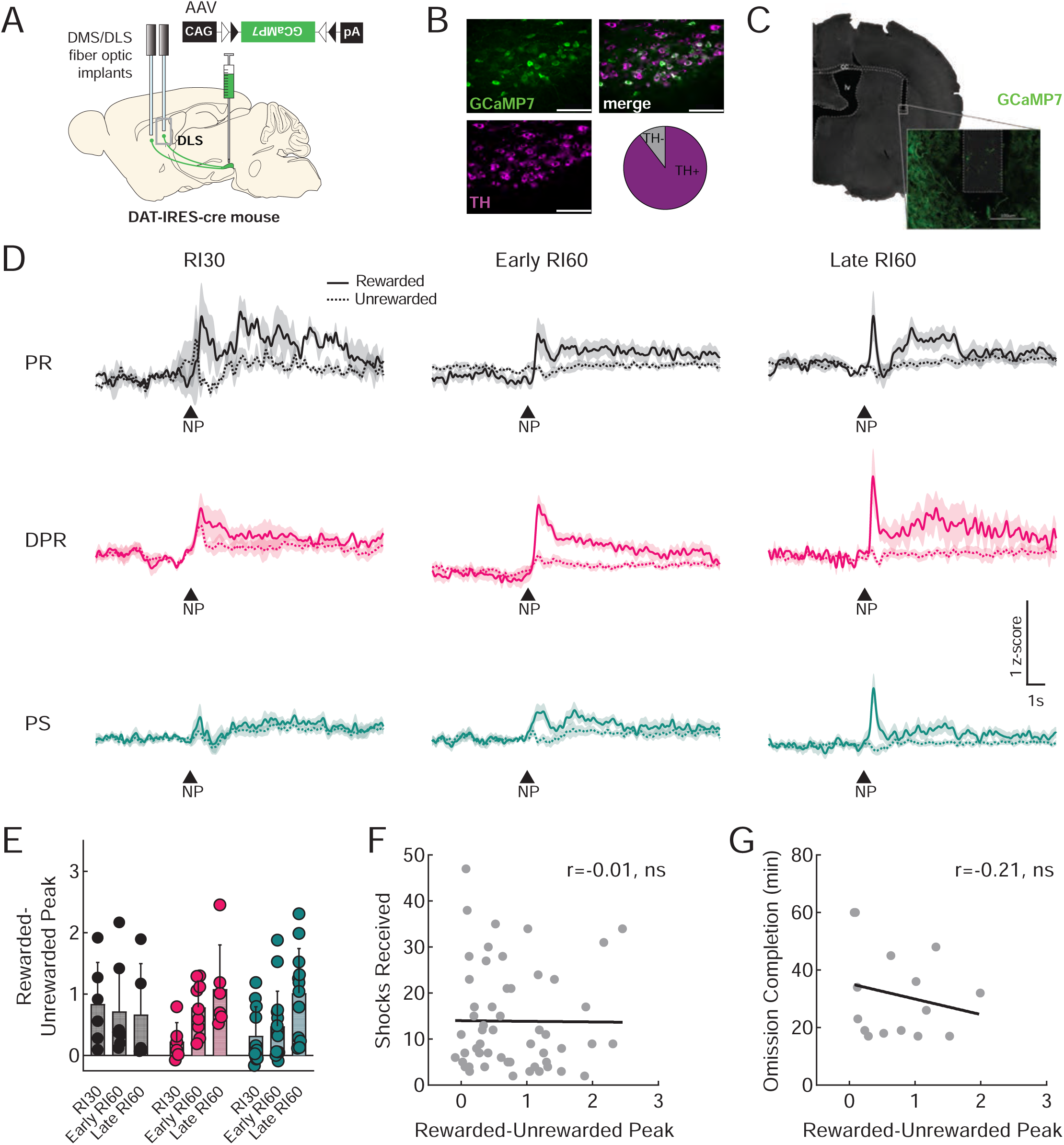
Dopamine signals in the DLS do not predict punishment-resistant or omission-resistant reward-seeking. **A.** Viral injection and probe placement strategy for recording from dopamine terminals in the DMS and DLS. DLS is highlighted here. **B.** High magnification (40x) images of SNc showing GCaMP7b expression in green, TH positive cells in magenta, and the merged image. Scale bars are 100μm. Quantification of GCaMP7b-expressing cells that are TH+ is shown; n=193 cells. **C.** Representative image showing probe placement in DLS. Area of magnification shows GCaMP7b expression in dopaminergic axons near the probe site in green. Scale bar is 100μm. cc=corpus callosum, lv=lateral ventricle. **D.** Peri-stimulus time histograms (PSTHs) showing the average signal from DLS dopamine terminals at the time of rewarded (solid) and unrewarded (dashed) nosepokes (NP) for each phenotype during RI30 training, early in training (RI60 day one or two), and late in training (RI60 day eleven or twelve). Shaded region represents SEM. Punishment resistant (PR; black), delayed punishment resistant (DPR; pink), or punishment sensitive (PS; teal). **E.** Quantification of the average “difference score” for DLS dopamine terminal signals observed in response to rewarded and unrewarded nosepokes (calculated as the peak at the time of rewarded nosepokes minus the peak at the time of unrewarded nosepokes). Error bars represent SD. **F.** Correlation of shocks received in shock probe sessions and “difference score” in DLS dopamine terminals (r=-0.01, ns). **G.** Correlation of time to complete omission and “difference score” in DLS dopamine terminals (r=-0.21, ns). See also Supplemental Figure 4.

To analyze these data, we first examined DMS dopamine axon activity occurring during rewarded and unrewarded nosepokes. We histologically verified that GCaMP was expressed in dopaminergic (TH+) neurons in the medial SNc and found that on average 82.86% of neurons expressing GCaMP also expressed TH (Fig. 3B). We confirmed that probe locations in the DMS were correct (Fig. 3C, S3A). Histology also verified robust GCaMP expression in dopaminergic axons in the DMS (Fig. 3C, inset). We compared DMS dopamine axon activity in RI60-trained mice (PR, DPR, PS; Fig. 3D) and in RR20-trained mice (RR20; Fig S3D) across training. Peaks in DMS dopamine axon activity at the time of a rewarded nosepoke were much clearer in PR and DPR mice than in PS or RR mice, an observation that was not true simply due to poor signal in PS or RR20 mice, as all mice included in the analysis had similar frequencies and amplitudes of GCaMP events across the entire training session (Fig. S3B-C). A main effect of training time on the frequency of all GCaMP events was observed, but was the same across all groups (Fig. S3B). Clear peaks in response to rewarded nosepokes were observed in PR mice even in the RI30 stage of training, whereas peaks in response to rewarded nosepokes emerged more slowly over the course of extended RI60 training in DPR mice (Fig. 3D). We also noted that mice varied in whether their DMS dopamine axons showed positive or negative responses to unrewarded nosepokes. In RI30 recordings, unrewarded nosepokes tended to give upward deflections in all groups, however, negative deflections appeared over the course of RI60 training, particularly in the PR and DPR groups. The combination of a positive peak in DMS dopamine axon activity for rewarded nosepokes and a negative peak for unrewarded nosepokes creates a notable difference in dopamine axon activity in response to the same motor action depending on the outcome, which is information that could be used to influence future behavior. We calculated a rewarded-unrewarded peak score for each mouse and compared whether the scores were altered by training stage or by behavioral phenotype (PR, DPR, or PS). There was a significant effect of training stage (Fig. 3E; mixed-effects analysis, F_1.61,29.8_=13.83, p<0.001), a significant effect of phenotype (F_2, 33_= 8.16, p<0.01) and a significant interaction between the two (F_4,37_= 3.29, p<0.05).

From these data, it appeared that the peaks in DMS dopamine axon activity tracked with the development of punishment-resistant behavior. We therefore wanted to assess whether an individual’s rewarded-unrewarded peak score (regardless of its classification as PR, DPR, or PS) could predict whether it would tolerate shocks in the shock probe sessions. Indeed, the shocks received were significantly correlated with the DMS rewarded-unrewarded peak score on a mouse-by-mouse basis (Fig. 3F; irrespective of our group classifications; r=0.53, p<0.0001). In contrast, performance on the omission probe was not correlated with this score (Fig. 3G; r=0.1, ns).

We also examined DMS dopamine axon signals surrounding the time of rewarded and unrewarded port entries and noticed ramping activity preceding the rewarded port entries (Fig. S3E-F). Such ramping towards reward has been described primarily in VTA-NAc dopamine circuits (Guru et al., 2020; Hamid et al., 2015; Howe et al., 2013; Kim et al., 2020; Mohebi et al., 2019), but has also been described in DMS dopamine axons in another recent pre-printed study (Hamid et al., 2019). Here, we additionally note that ramping in DMS dopamine axons is more prominent in PR and DPR mice than in PS or RR20 mice (Fig. S3E-F). To encapsulate ramping activity quantitatively, we measured the area under the curve (AUC) of our fiber photometry signal from -5 to 0s relative to the rewarded port entry for each mouse. The AUC was variable, but notably the DPR group of mice showed a significant increase from early RI60 to late RI60 training (Fig. S3F; mixed-effects analysis, interaction of time and phenotype, F_3,19_= 4.77, p<0.05, Bonferroni, p<0.05).

### Dopamine signals in the DLS do not predict punishment-resistant or omission-resistant reward-seeking

In recordings from the same mice, we next examined DLS dopamine axon activity occurring during nosepokes to see if it differed from activity in the DMS (Fig. 4A). We verified the expression of virus in SNc (Fig. 4B; 89.56% of GCaMP neurons also expressed TH) and the probe placements in DLS (Fig. 4C, S4A). We also noted that the DLS signals in all groups of mice had similar frequencies and amplitudes of GCaMP events (Fig. S4B-C). DLS dopamine axon signals at the time of a rewarded nosepoke differed from the DMS dopamine axon signals in that they had both an immediate component and a prolonged, delayed component (Fig. 4D, S4D). DLS dopamine axon signals following an unrewarded nosepoke were variable. During RI30, there tended to be positive deflections in the signal for unrewarded nosepokes, however, later in training negative deflections emerged. To test whether these DLS dopamine axon signals in response to nosepokes bore any relationship to the punishment-resistant phenotype of the mice, we calculated a rewarded-unrewarded peak score as we had for the DMS dopamine axon signals. A mixed-effects analysis revealed a significant effect of training stage (mixed-effects analysis, F_2,35_= 3.53, p<0.05), indicating that the signals do change over the course of training, but no significant effect of group. For completeness, we also looked at whether these signals correlated on an individual basis with shocks received in the shock probe or omission completion time and found no correlations (Fig. 4F-G; r=-0.01, ns and r=-0.21, ns, respectively).

We also assessed whether the prolonged elevation of DLS dopamine axon activity after a rewarded nosepoke, which was often sustained for 10 seconds or more, was related to the development of punishment- or omission-resistance. We quantified the prolonged activity as the area under the curve (AUC) from 2-10 seconds after the nosepoke. AUC did not correlate with the performance of individuals on the shock or omission probes (Fig. S4E-F) nor did it differ statistically across groups or training time (Fig. S4G).

We were surprised that we could not observe a correlation of DLS dopamine axon signals with behavior, despite the fact that they changed over time. To get another view of these signals, we tried grouping mice by their performance on the omission probe rather than the shock probe. However, we saw no obvious differences in the development of dopamine axon signals in either the DMS or the DLS depending on omission probe performance (Fig. S4H).

Finally, we also looked at DLS dopamine axon signals aligned to the time of port entries (Fig. S4I). Here, we observed very weak ramping prior to a rewarded port entry, with the peak of the signal occurring after port entry. Quantification of the AUC of our fiber photometry signal from -5s to 0s relative to the rewarded port entry did not show evidence that pre-port entry activity tracked with the behavioral phenotype (Fig. S4J). The lack of ramping in DLS dopamine axons in comparison to DMS dopamine axons we observed is notably similar to what was observed in another recent study, even though the imaging methods and behaviors used in that study are distinct from ours (Hamid et al., 2019). The functional significance of this difference in ramping activity between the DMS and DLS dopamine axons will be an important area for future investigation.

### Optogenetic excitation of dopamine terminals in DMS at the time of a rewarded nosepoke accelerates the development of punishment-resistant reward-seeking

Since peaks in DMS dopamine axon activity in response to rewarded nosepokes predicted the development of punishment-resistant reward-seeking, we decided to test if stimulation of DMS dopamine axons at the time of a rewarded nosepoke could cause punishment-resistant behavior to emerge. We used the excitatory opsin ChR2 to stimulate DMS dopamine axons. An AAV expressing cre-dependent ChR2 (AAV5-EF1α-DIO-hChR2(H134R)-EYFP) was injected into the SNc of DAT-IRES-cre mice to express ChR2 specifically in dopamine neurons. A fiber optic probe was placed above the DMS to allow light stimulation of dopamine terminals only within the DMS (Fig. 5A). Previous work has shown that DMS-projecting dopamine neurons have minimal collateralization outside the DMS (Lerner et al., 2015), so this stimulation should be specific even if back-propagating action potentials are generated. We verified that this strategy led to the expression of ChR2 and EYFP in dopamine (TH+) neurons (Fig. 5B-C) and that fiber optics were appropriately placed in the DMS, matching the coordinates used for fiber photometry recordings (Fig. S5A).

**Figure 5.**
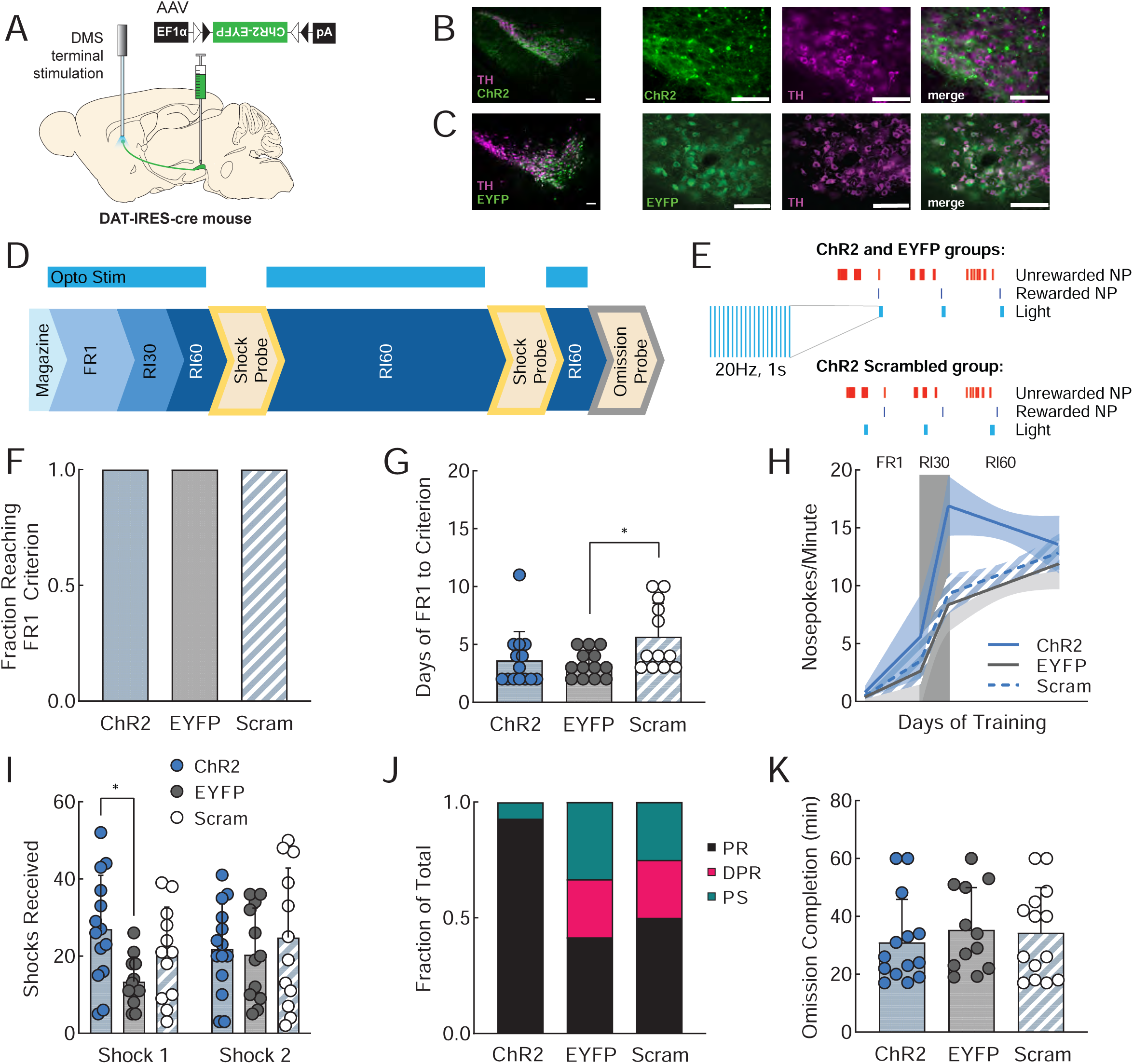
Optogenetic excitation of dopamine terminals in DMS at the time of a rewarded nosepoke accelerates the development of punishment-resistant reward-seeking. **A.** Viral injection and probe placement strategy for stimulation of dopamine terminals in the DMS. **B.** Low (10x) and high (40x) magnification images of SNc showing ChR2-EYFP expression in green, TH positive cells in magenta, and the merged image. Scale bars are 100μm. **C.** Low (10x) and high (40x) magnification images of SNc showing EYFP expression in green, TH positive cells in magenta, and the merged image. Scale bars are 100μm. **D.** Training timeline showing sessions during which optogenetic stimulation was delivered (FR1, RI30, RI60). **E.** Schematic of stimulation parameters. A 1s, 20Hz burst of stimulation was paired with rewarded nosepokes for ChR2 and EYFP groups and the same stimulation was paired with a random subset of nosepokes for ChR2 Scrambled animals. **F.** Fraction of mice in each group that reached the criterion to move on from FR1 training (see methods). ChR2 (blue; n=14), EYFP (gray; n=13), ChR2 scrambled (blue stripe; n=12). **G.** Average days required for animals in each group to reach criterion to move on from FR1. Error bars represent SD *p<0.05. **H.** Segmental linear regression showing the slope of nosepokes made per minute in FR1, RI30, and RI60 schedules. Shaded region represents 95% confidence bands. **I**. Average shocks received on early and late shock probes for each group. Error bars represent SD *p<0.05. **J.** Fraction of each behavioral phenotype (punishment resistant=black, delayed punishment resistant=pink, punishment sensitive=teal) in each group. **K.** Average time to complete the omission probe (earn 50 rewards; max 60 minutes) for each group. Errors bars represent SD. See also Supplemental Figure 5.

Beginning with FR1 training, ChR2 animals received 1s, 20Hz trains of light stimulation on every rewarded nosepoke (Fig. 5D-E). Importantly, stimulation of DMS dopamine terminals was given during FR1/RI30/RI60 training but not during shock probes when punishment-resistant reward seeking was assessed. Therefore, any effects of the stimulation on probe performance are not due to acute effects of dopamine terminal stimulation, but are caused by differences in learning during the training sessions. In addition to animals expressing ChR2 and receiving stimulation on rewarded nosepokes, there were two control groups: EYFP controls that received a fluorophore-only control virus injected into the SNc (AAV5-EF1α-DIO-EYFP) and the same pattern of light stimulation, and “scrambled” controls that received the ChR2 virus, but had their DMS dopamine terminals stimulated on a random subset of nosepokes rather than on rewarded nosepokes. The scrambled control was important as these mice received the same amount of dopamine terminal stimulation as the ChR2 group and this stimulation reinforced the same action (a nosepoke). Therefore, the only difference between the ChR2 and ChR2 Scrambled groups was whether or not the dopamine terminal stimulation they received boosted vs degraded the ability of the natural DMS dopamine signal to differentiate between externally rewarded and unrewarded actions.

All three groups (ChR2, EYFP, and ChR2 Scrambled) learned the FR1 task and were advanced to the RI schedules (Fig. 5F). However, mice in the ChR2 Scrambled group took significantly longer to perform to criterion on the FR1, perhaps indicating that scrambled stimulation caused an initial learning impairment (Fig. 5G; unpaired t-test, p<0.05). After acquiring FR1, however, all mice performed the RI60 task and received approximately the same number of rewards per minute and per session (Fig. S5B-C). ChR2 mice escalated their nosepoking much faster than the two groups of control mice during early training (FR1 and RI30), then leveled or even dropped their nosepoking rates over the course of extended RI60 training (Fig 5H). When tested on the first shock probe after minimal RI60 training, ChR2 mice were significantly more resistant to punishment than EYFP mice (Fig. 5I; Tukey’s multiple comparison, p<0.05). This difference faded with extended training as punishment-resistant behavior also began to emerge naturally in the control groups. We categorized mice from this experiment as PR, DPR and PS using our previously defined criteria. We found that ChR2 mice were extremely likely to be categorized as PR, whereas the EYFP and ChR2 Scrambled mice were distributed as expected across groups (Fig. 5J). Notably, under ChR2 stimulation, 100% of male mice were PR mice (Fig. S5D). A large majority of female mice (∼71%) were also categorized as PR – despite the fact they were unlikely to be PR in our GCaMP, EYFP, and ChR2 Scrambled groups – indicating that DMS dopamine terminal stimulation on rewarded nosepokes can drive both sexes to quickly develop punishment-resistant reward-seeking (Fig. S5D, cf. Figs. 2A, S2A).

We also looked to see whether DMS dopamine terminal stimulation influenced performance on a final omission test after extended RI60 training. It did not. ChR2, EYFP, and ChR2 Scrambled mice all took the same amount of time to complete the omission probe (Fig. 5K). These data confirm that the correlations we observed between DMS dopamine axon activity and the development of punishment-resistant behavior are causal, and specific to the development of punishment-resistant behavior. The fact that DMS dopamine terminal stimulation did not affect omission-resistance suggests that these two forms of inflexible behavior can be supported by distinct neural circuits.

### Optogenetic inhibition of dopamine terminals in DMS interferes with action-outcome learning

Promoting DMS dopamine activity in response to rewarded nosepokes accelerated the development of punishment-resistant reward-seeking. We therefore wanted to ask the opposite question: would inhibiting DMS dopamine activity delay its development? For this experiment, we performed bilateral inhibition of DMS dopamine axons using the inhibitory opsin eNpHR3.0 (Gradinaru et al., 2010). An AAV expressing cre-dependent NpHR (AAV5-EF1α-DIO-eNpHR3.0-EYFP) or a fluorophore-only control virus (AAV5-EF1α-DIO-EYFP) was injected into the SNc of DAT-IRES-cre mice to express NpHR (or EYFP) specifically in dopamine neurons. Fiber optic probes were placed above the DMS to allow light delivery (Fig. 6A). We verified that this strategy led to the expression of NpHR and EYFP in dopamine (TH+) neurons (Fig. 6B-C) and that fiber optics were appropriately placed in the DMS (Fig. S6A).

**Figure 6.**
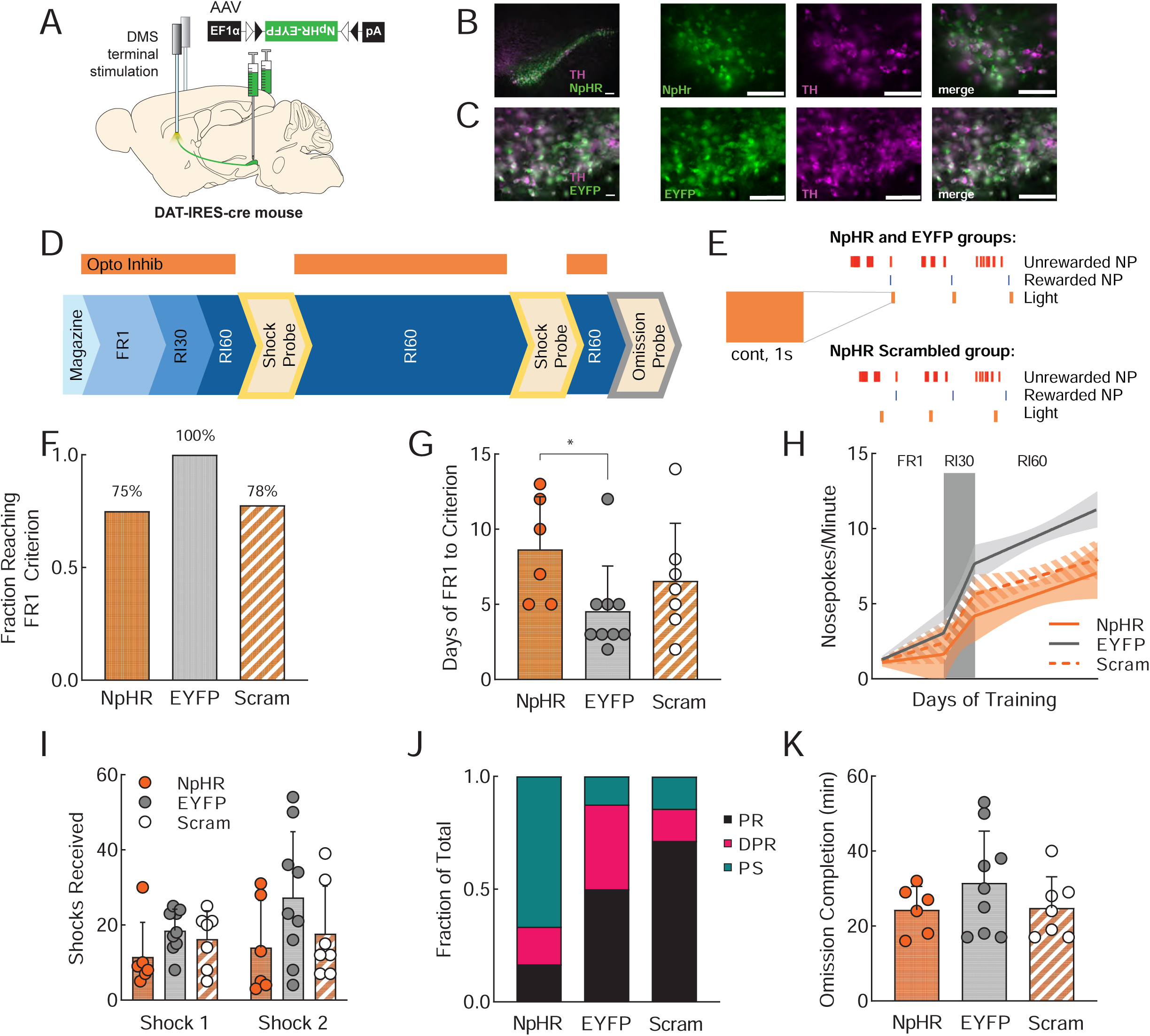
Optogenetic inhibition of dopamine terminals in DMS interferes with action-outcome learning. **A.** Viral injection and probe placement strategy for inhibition of dopamine terminals in bilateral DMS. **B.** Low (10x) and high magnification (40x) images of SNc showing NpHR-EYFP expression in green, TH positive cells in magenta, and the merged image. Scale bars are 100μm. **C.** Low (10x) and high magnification (40x) images of SNc showing EYFP expression in green, TH positive cells in magenta, and the merged image. Scale bars are 100μm. **D.** Training timeline showing sessions during which optogenetic stimulation was delivered (FR1, RI30, RI60). **E.** Schematic of stimulation parameters. A 1s continuous light delivery was paired with rewarded nosepokes for NpHR and EYFP groups and the same light was paired with a random subset of nosepokes for NpHR Scrambled animals. **F.** Fraction of mice in each group that reached the criterion to move on from FR1 training (see methods). NpHR (orange; n=6 of 8), EYFP (gray; n=9), NpHR scrambled (orange stripe; n=7 of 9). **G.** Average days required for animals in each group to reach criterion to move on from FR1 (for those that did). Error bars represent SD *p<0.05. **H.** Segmental linear regression showing the slope of nosepokes made per minute in FR1, RI30, and RI60 schedules. Shaded region represents 95% confidence bands. **I**. Average shocks received on early and late shock probes for each group. Error bars represent SD. **J.** Fraction of each behavioral phenotype (punishment resistant=black, delayed punishment resistant=pink, punishment sensitive=teal) in each group. **K.** Average time to complete the omission probe (earn 50 rewards; max 60 minutes) for each group. Errors bars represent SD. See also Supplemental Figure 6.

Mice were divided into three groups, as for the stimulation experiment. NpHR mice received a 1s continuous pulse of light on every rewarded nosepoke (Fig. 6D-E). EYFP mice received the same light stimulation, but lacked NpHR. NpHR Scrambled mice received a 1s continuous pulse of light on a random subset of nosepokes. We chose to begin the light delivery on FR1 to parallel the design of the excitatory optogenetics experiment (Fig. 5), however, DMS dopamine terminal inhibition resulted in a learning deficit. In other experiments, 100% of mice had quickly reached criterion for FR1 performance, showing that they were able to learn the association of their action (nosepoke) with an outcome (delivery of a pellet). However, only 75% of NpHR mice and 78% of NpHR Scrambled mice reached criterion (Fig. 6F). The other 25% and 22% of mice, respectively, were dropped from the study after more than 14 days (mean+2SD) of unsuccessful FR1 training. Of the mice that did pass our FR1 criterion, the NpHR mice required more days of training than EYFP control mice to reach FR1 criterion (Fig. 6G; unpaired t-test, p<0.05). Both of the groups receiving inhibition of their DMS dopamine terminals also displayed a reduced escalation in nosepoke rates over training (Fig. 6H). On a day-by-day basis, all groups received rewards at approximately the same rate (Fig. S6B), although when pooled over days of RI60, NpHR mice received slightly fewer rewards than EYFP mice (Fig. S6C; p<0.05). Overall, we conclude that optogenetic inhibition of dopamine terminals in DMS at the time of the nosepoking action interferes with action-outcome learning in a manner that is not specific to the time of the rewarded nosepoke.

For the mice that passed FR1 and were able to complete the experiment, we tested punishment-resistance on the shock probe both early and late in training. We did not observe significant differences between groups in the total number of shocks received (Fig. 6I). However, when we sorted the mice into PR, DPR and PS groups, we noted that the NpHR group had an increased incidence of PS mice, while the NpHR Scrambled group had an increased incidence of PR mice, compared to EYFP controls (Fig. 6J). The effects of NpHR inhibition on PR/DPR/PS phenotype are driven by stark effects in male mice (Fig. S6D). We also tested these mice on the omission probe at the end of training, but did not observe significant differences in omission time (Fig. 6K). These data suggest that after initial learning delays due to DMS dopamine terminal inhibition are overcome, inhibition specifically on rewarded nosepokes may also delay the development of punishment-resistance, particularly in male mice.

### Optogenetic excitation of dopamine terminals in DLS at the time of a rewarded nosepoke does not influence instrumental learning or behavioral flexibility

Peaks in DMS dopamine axon activity in response to rewarded nosepokes predicted the development of punishment-resistant reward-seeking (Fig. 3F), but peaks in DLS dopamine axon activity did not (Fig. 4F). Therefore, we hypothesized that the stimulation of DLS dopamine terminals following rewarded nosepokes would not affect the development of punishment-resistant reward-seeking. Similarly, since peaks in DLS dopamine axon activity did not correlate with omission completion time, we hypothesized that DLS dopamine terminal stimulation would not affect omission completion time. Nevertheless, we decided to test the effects of DLS dopamine terminal stimulation as a counterpoint to the effects of DMS dopamine terminal stimulation (Fig. 5, S5). In other words, can stimulating *any* dopamine signal boost the development of punishment resistance, or is this effect specific to DMS?

We performed the same experiment as in Figure 5, but targeting DLS instead of DMS. An AAV expressing cre-dependent ChR2 (AAV5-EF1α-DIO-hChR2(H134R)-EYFP) or a fluorophore-only control virus (AAV5-EF1α-DIO-EYFP) was injected into the SNc of DAT-IRES-cre mice to express ChR2 or EYFP specifically in dopamine neurons. A fiber optic probe was placed above the DLS to allow light stimulation of dopamine terminals (Fig. 7A). Fiber placements and appropriate ChR2 and EYFP expression were verified by histology (Fig. 7B-C, S7A). Light stimulation (1s, 20Hz) was delivered during training sessions, beginning with FR1 (Fig. 7D-E). All mice in this experiment quickly reached FR1 criterion and advanced to RI training (Fig. 7F-G). However, in contrast to the DMS dopamine stimulation experiment, all groups of mice (ChR2, EYFP, and ChR2 Scrambled) behaved similarly. All groups of mice escalated their nosepoking at the same rates (Fig. 7H). All received similar numbers of shocks in both of the shock probes (Fig. 7I) and had similar distributions of PR, DPR, and PS mice (Fig. 7J). No significant differences were observed in omission completion time at the end of the experiment (Fig. 7K). We conclude that DLS dopamine terminal stimulation immediately following a rewarded nosepoke does not influence the development of punishment-resistant or omission-resistant reward-seeking behavior. However, the behavioral consequence of the more prolonged/delayed activity of DLS dopamine axons we observed following rewarded nosepokes (Fig. 4D) remains to be explored in future experiments.

**Figure 7.**
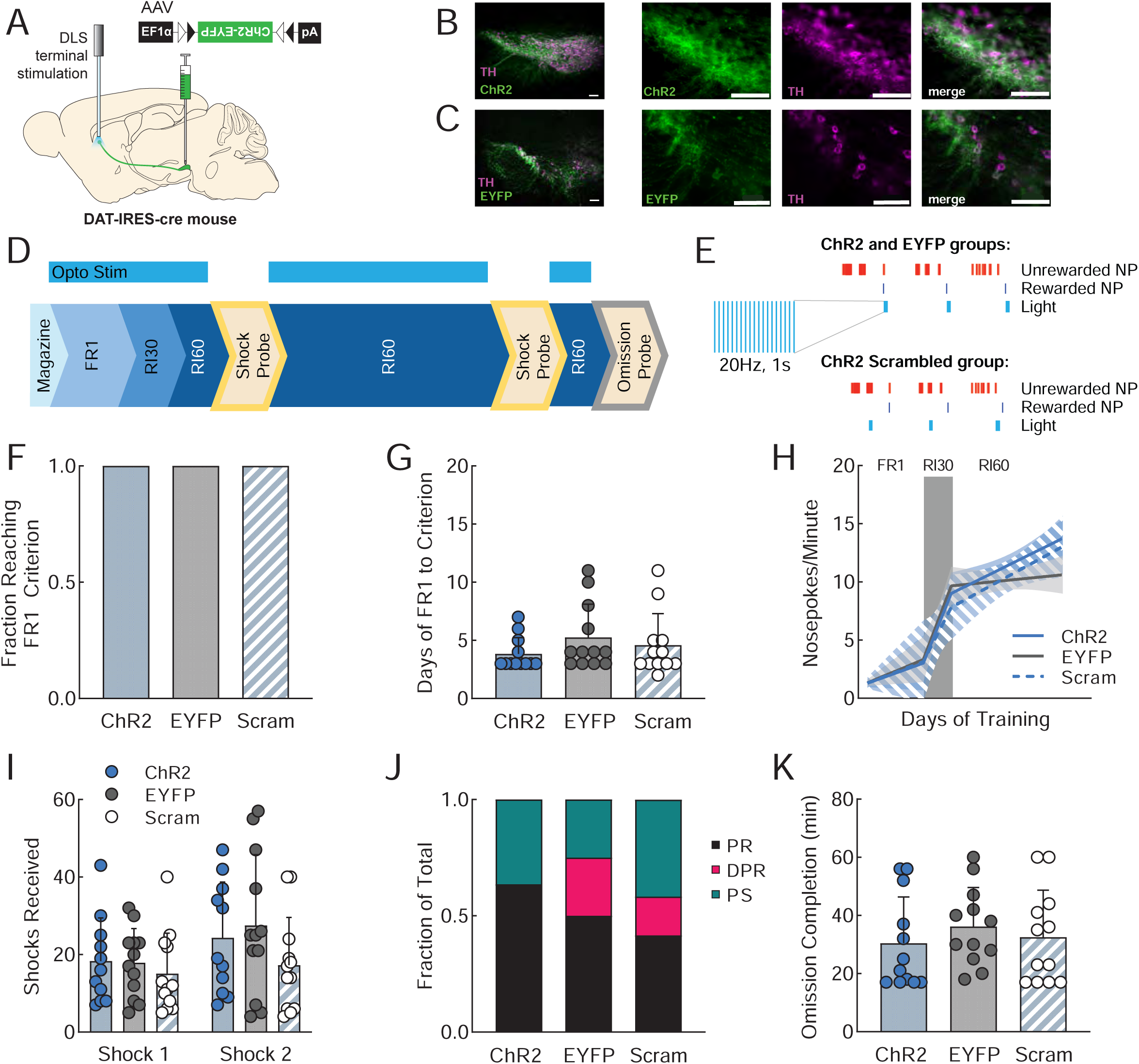
Optogenetic excitation of dopamine terminals in DLS at the time of a rewarded nosepoke does not influence instrumental learning or behavioral flexibility. **A.** Viral injection and probe placement strategy for stimulation of dopamine terminals in the DLS. **B.** Low (10x) and high (40x) magnification images of SNc showing ChR2-EYFP expression in green, TH positive cells in magenta, and the merged image. Scale bars are 100μm. **C.** Low (10x) and high (40x) magnification images of SNc showing EYFP expression in green, TH positive cells in magenta, and the merged image. Scale bars are 100μm. **D.** Training timeline showing sessions during which optogenetic stimulation was delivered (FR1, RI30, RI60). **E.** Schematic of stimulation parameters. A 1s, 20Hz burst of stimulation was paired with rewarded nosepokes for ChR2 and EYFP groups and the same stimulation was paired with a random subset of nosepokes for ChR2 Scrambled animals. **F.** Fraction of mice in each group that reached the criterion to move on from FR1 training (see methods). ChR2 (blue; n=11), EYFP (gray; n=12), ChR2 scrambled (blue stripe; n=12). **G.** Average days required for animals in each group to reach criterion to move on from FR1. Error bars represent SD. **H.** Segmental linear regression showing the slope of nosepokes made per minute in FR1, RI30, and RI60 schedules. Shaded region represents 95% confidence bands. **I**. Average shocks received on early and late shock probes for each group. Error bars represent SD. **J.** Fraction of each behavioral phenotype (punishment resistant=black, delayed punishment resistant=pink, punishment sensitive=teal) in each group. **K.** Average time to complete the omission probe (earn 50 rewards; max 60 minutes) for each group. Errors bars represent SD. See also Supplemental Figure 7.

## DISCUSSION

Compulsive behavior is a defining feature of disorders such as substance use disorder (SUD) and obsessive-compulsive disorder (OCD). There is some evidence suggesting that corticostriatal circuits control the expression of established compulsions, but little is known about the mechanisms regulating the development of compulsions (Lüscher & Janak, 2021). We hypothesized that dopamine – a key neuromodulator regulating corticostriatal synaptic plasticity – could play a role in sculpting the emergence of compulsive behavior, defined as punishment-resistant reward-seeking. However, it was unclear whether the dopamine signals most relevant punishment-resistant reward-seeking would lie in the DMS, which is linked to action-outcome learning, or the DLS, which is linked to habit formation. We addressed this question by using dual-site fiber photometry to record the activity of dopamine axons in the dorsomedial striatum (DMS) and dorsolateral striatum (DLS) during a task (RI60) that promotes punishment-resistant reward-seeking. We identified task-relevant increases in DMS dopamine axon activity as a key feature of neural circuit activity predicting which individual animals will develop punishment-resistance. Specifically, DMS dopamine axon activity that effectively discriminated between rewarded and unrewarded nosepokes was correlated with the development of punishment-resistant reward-seeking. Therefore, we hypothesized that it is the discrimination of rewarded vs unrewarded actions by DMS dopamine – not the general dopamine-mediated reinforcement of the nosepoking action – that drives compulsion. To test this hypothesis, we optogenetically stimulated dopamine terminals in the DMS during learning. By creating peaks of DMS dopamine axon activity on rewarded nosepokes, we accelerated the development of punishment-resistant reward-seeking without influencing another form of inflexible responding (omission) more closely related to habit formation. Stimulating DMS dopamine terminals on random nosepokes did not affect the behavior, indicating that the timing of DMS dopamine stimulation with respect to external outcomes was important. Stimulating DLS dopamine terminals on rewarded nosepokes did not accelerate the development of punishment-resistant reward-seeking. These results support a model in which a properly timed DMS dopamine signal is specifically linked to the emergence of punishment-resistant reward-seeking.

The finding that DMS dopamine signaling promotes the development of punishment-resistant reward-seeking fits well with observations indicating that OFC inputs to the DMS are important for this behavior. Punishment-resistance in a paradigm in which animals self-stimulate their VTA dopamine neurons depends on enhanced excitability of OFC and potentiation of the OFC-DMS pathway (Pascoli et al., 2015, 2018). Increased OFC-DMS activity has also been associated with punishment-resistant methamphetamine-seeking (Hu et al., 2019b). Further, the OFC-DMS pathway is strengthened by repeated non-contingent injections of cocaine (Bariselli et al., 2020), and OFC neurons represent cocaine preference in cocaine-preferring rats (Guillem & Ahmed, 2018), indicating that the OFC-DMS circuit is one that could be co-opted by addictive drugs to provoke compulsive use. Our results do not directly test inputs from OFC. However, together with these previous findings, they suggest that an over-strengthening of the OFC-DMS pathway, perhaps by dopamine-dependent synaptic plasticity mechanisms, could promote punishment-resistance on a goal-directed rather than a habitual basis (Gremel et al., 2016; Gremel & Costa, 2013; Hogarth, 2020; Lüscher et al., 2020).

Although some previous studies have observed a progression from habit to compulsion after extended training (Everitt & Robbins, 2005, 2016; Giuliano et al., 2019) we suggest that these observations could be due to a common upstream driver of DMS and DLS function, rather than a direct and necessary link between habit formation and the development of compulsive behavior. In our experiments, extended RI60 training led to habit formation (as it has been previously documented to do; Derusso et al., 2010; Yin et al., 2006) in addition to compulsion. Thus, these two endpoints could easily be confused. However, by analyzing individual differences in behavior, we determined that the development of habits and compulsions do not inevitably develop together, consistent with the findings of Singer et al. and others (2017; Olmstead et al., 2001). Nevertheless, our results do not rule out the possibility that there are both DMS- and DLS-dependent routes to developing punishment-resistance, which could be invoked under different circumstances (e.g. different training schedules, or different modalities of reward).

### Unanswered Questions

Further studies are needed on several topics. First, we need to more explicitly examine the relationship between DLS dopamine signals and habit. Previous approaches using lesions and pharmacology are suggestive of a relationship, but do not provide temporal specificity. Here, we observed novel temporal dynamics in the DLS dopamine signal, which differentiate it from the signal in the DMS. The DLS dopamine signal has an immediate peak following a rewarded nosepoke as well as prolonged activity above baseline. We did not find strong correlations of this signal with individual behavior, but alternative tests for habit, such as outcome devaluation tests, may reveal a stronger association in future studies. DLS dopamine may also be involved in other tasks. For example, one recent study linked high levels of extracellular dopamine in the DLS with high impulsivity in a delay-discounting task (Moreno et al., 2021). Another study linked a molecularly-defined population of dopamine neurons that primarily projects to the DLS (Aldh1a1+ dopamine neurons) to motor learning on the accelerating rotarod (Wu et al., 2019). These examples highlight how much more parameter space there is still to be explored in terms of the relationship between DLS dopamine and behavior.

Second, future work should examine temporal patterns. As a control for our optogenetics experiment, we included a group of mice that received DMS dopamine terminal stimulation on random nosepokes. Creating peaks in DMS dopamine on random nosepokes did not have the same effect as creating these peaks on rewarded nosepokes. The random stimulation data indicate that the temporal pattern of dopamine activity in the DMS matters, but it remains to be determined *why* the pattern is important. For example, future studies could examine whether the pattern of cortical inputs to the DMS is different during rewarded vs unrewarded nosepokes. If different cortical inputs to the DMS are active during rewarded vs unrewarded nosepokes, dopamine release at these distinct times would reinforce the strength of different corticostriatal synapses.

Third, further studies could amplify our understanding of the DMS mechanism we observed, and clarify how it relates to the finding that projections from the lateral hypothalamus (LH) to the VTA could bidirectionally control compulsive sucrose seeking as in a previous study (Nieh et al., 2015). We do not know how this LH-VTA circuit might interact with DMS dopamine signaling; however, there are several possibilities. Most simply, some VTA dopamine neurons project to the DMS (Beier et al., 2015; M. W. Howe & Dombeck, 2016). Additionally, DMS-projecting dopamine neurons receive inputs from LH, VTA and NAc, any of which could easily interconnect the circuitry (Lerner et al., 2015).

### Larger Implications

To understand behavior, we need to grapple with individual differences. Not all mice (or people) who try drugs become compulsive users. Large individual variability in compulsivity has been observed in animals working for drugs such as cocaine and alcohol (Giuliano et al., 2019; Siciliano et al., 2019). Our findings underline the necessity of this approach; we saw large individual differences as mice worked for natural rewards. We identified one reason for this variability: the different strategies used by individual animals to deal with uncertainty in the RI60 schedule. In our study, some mice maximized their rate of reward retrieval by nosepoking at high rates, while other mice conserved effort. This finding also sheds light on the poorly understood question of how punishment-resistant reward-seeking emerges over time. In addition to differences in neural signals, we also observed different behavioral strategies prior to the experience of punishment, which allowed us to predict which animals will go on to become punishment-resistant. This pattern suggests there is a predisposition to towards developing punishment-resistance present in individuals before they confront punishments, rather than a stochastic process occurring during the experience of punishment, to generate punishment-resistance.

One source of individual variability in our studies was sex: male mice were more likely to be punishment-resistant than females. Nevertheless, sex could not fully explain individual variability. Furthermore, the correlation between DMS dopamine axon signaling and the development of punishment-resistance was not sex-dependent, and DMS dopamine terminal stimulation could induce both sexes to transition to punishment-resistance. We therefore suspect that sex differences influencing differences in the likelihood of males and females to develop punishment resistance occur upstream of dopamine neurons.

Understanding compulsive behavior in humans and rodents also requires understanding whether or not there is variability associated with the nature of the rewarding substance. In this study, we examined the behavior of mice during learning to pursue sucrose rewards. Compulsive sucrose-seeking is relatively understudied; most previous studies have examined compulsive drug-seeking. It is important to understand behavior across natural and manufactured rewards of various kinds. By studying compulsive seeking of a natural reward, we can better understand the evolutionary context under which this behavior developed, perhaps working to promote what might be called “grit” in the face of life’s inevitable challenges. Furthermore, understanding how compulsive drug-seeking and compulsive sucrose-seeking relate to each other can elucidate how concepts from SUD should be applied to our understanding other disorders like eating disorders, gambling disorders, or OCD (among others). While the circuit mechanism promoting compulsive sucrose-seeking we have identified is exciting, we need to ascertain whether it would similarly drive the development of compulsive drug-seeking. Conversely, previously identified circuits for compulsive drug-seeking might or might not impact compulsive sucrose-seeking. In fact, one behavioral study directly examined whether the development of compulsive drug-seeking and compulsive sucrose-seeking were correlated and found that they were NOT: animals that became compulsive for one type of reward did not necessarily become compulsive for the other type (Datta et al., 2018). Thus, it is imperative to examine in more detail whether all rewards do or do not activate the same circuits for compulsivity.

In summary, we have identified DMS dopamine signaling as a key part of the circuitry that drives the emergence of compulsive behavior in the context of natural reward-seeking. The data presented here set the stage for interesting new studies in a variety of areas. Examining how the mechanisms we have identified contribute to the etiology of disorders such as SUD and OCD is of particular importance for translational impact.

## ACKNOWLEDGEMENTS

We thank members of the Lerner laboratory for helpful discussions and critical feedback on the manuscript. We thank G. Palissery and M. Holla for assistance with surgeries, histology, data collection and entry. pGP-AAV-CAG-FLEX-jGCaMP7b-WPRE was a gift from Douglas Kim & the GENIE Project. AAV5-EF1α-DIO-hChR2(H134R)-EYFP was a gift from Karl Deisseroth. This work was supported by an NIH K99/R00 Award (R00MH109569) and a NARSAD Young Investigator Award from the Brain & Behavior Research Foundation to T.N.L., and an NIH Diversity Supplement (R00MH109569-04S1) to support C.V.C.

## AUTHOR CONTRIBUTIONS

Conceptualization, Methodology, Software, Validation, and Project Administration, J.L.S., C.V.C., and T.N.L.; Investigation, J.L.S., C.V.C., and A.S.B.; Data Curation and Visualization, J.L.S., J.M.B., A.S.B., and T.N.L.; Formal Analysis, J.L.S., V.N.S. and J.M.B..; Writing - Original Draft, J.L.S.; Resources, Writing - Review & Editing, Supervision, and Funding Acquisition, T.N.L.

## STAR METHODS

### Lead Contact and Materials Availability

Further information and requests for resources and reagents should be directed to and will be fulfilled by the Lead Contact, Talia Lerner. This study did not generate new unique reagents.

### Experimental Model and Subject Details

#### Mice

Male and female *WT* (C57BL/6J) and (DAT)::IRES-Cre knockin mice (JAX006660) were obtained from The Jackson Laboratory and crossed in house. Only heterozygote transgenic mice, obtained by backcrossing to C57BL/6J wildtypes, were used for experiments. Littermates of the same sex were randomly assigned to experimental groups (fiber photometry-14 males, 22 females; DMS excitatory optogenetics-20 males, 19 females; DMS inhibitory optogenetics-13 males, 13 females; DLS excitatory optogenetics-18 males, 18 females). Adult mice at least 10 weeks of age were used in all experiments. Mice were group housed under a conventional 12h light cycle (dark from 7:00pm to 7:00am) with *ad libitum* access to food and water prior to operant training. All experiments were approved by the Northwestern University Institutional Animal Care and Use Committee.

### Method Details

#### Operant Behavior

Mice were food restricted to 85% of ad libitum body weight for the duration of operant training. Mice were given one day of habituation to operant chambers (Med Associates) and tethering with patch cords (Doric Lenses) for one hour. They were then trained to retrieve food rewards (45 mg purified pellet, Bio-Serv) from a magazine port. For this magazine training, pellets were delivered to the port on a random interval (RI60) schedule non-contingently for one hour. Next, operant training began, with all training sessions lasting one hour or until 50 rewards had been earned. Mice were trained to associate nosepoking with reward on a fixed ratio (FR1) schedule where both nosepokes delivered a reward. They had to retrieve the reward (as measured by making a port entry following a rewarded nosepoke) before they could earn the next reward. After a mouse showed a preference for one nosepoke (>25 rewards on that side; average of 3.06 days), they were trained on FR1 on their preferred side only, with nosepokes on the other side having no consequence, until they received >30 rewards for a minimum of two consecutive days (average of 5.87 days). Mice that did not reached this criterion after 14 days of FR1 training (mean+2 SD), were removed from the study. Mice passing the FR1 criterion were then moved to either a random interval (n=36) or random ratio (n=7) schedule of reinforcement. Mice on the random interval schedule were trained on RI30 until they earned >30 rewards in one hour (average of 2.33 days), and then trained on RI60. Mice on a random ratio schedule of reinforcement were trained on RR10 until they earned >30 rewards in one hour (average of 2.71 days), and then trained on RR20 (Fig. 1A). For random interval and random ratio schedules, a normal distribution centered around the number indicated in the name of the schedule was used to create the schedule. The range for RI30 was from 15-45s, RI60 from 30-90s, RR10 from 6-14 nosepokes, and RR20 from 14-28 nosepokes.

#### Shock Probe

Mice were subjected to a footshock probe early and late in training (Fig. 1B) to evaluate their levels of punishment-resistance reward-seeking. These probes were performed under an FR1 schedule of reinforcement where a mild footshock (0.2mA, 1s) was paired with a subset of rewarded nosepokes on a RR3 schedule, so that, on average, every third rewarded nosepoke was paired with a footshock. During shock probes, the session ended after 60 minutes or a mouse was inactive (no nosepokes on the rewarded side) for >10 minutes. There was no maximum number of rewards.

#### Omission Probe

A subset of mice (n=20) were returned to RI60/RR20 training after the late footshock probe until their nosepoke rates returned to pre-shock levels. They then received a single omission probe session where they had to withhold nosepoking for 20 seconds in order to receive a single reward pellet. A nosepoke reset the 20 second timer. Each session ended after a mouse received 50 rewards or 60 minutes had elapsed.

#### Fear Conditioning

A trace fear conditioning paradigm, adapted from Lugo, Smith, and Holly (2014) was used in a naïve cohort of wild-type mice (n=13) to verify that our shock intensity (0.2mA) is aversive to the mice (Lugo et al., 2014). Mice were randomly assigned to cued or non-cued groups. On the first day, mice received 12 tone only or tone-shock pairings (2900 Hz tone) in a standard operant chamber (Med Associates). The next day, mice were placed in a different context (using white walls, white plastic flooring, and vanilla scent) and 12 tones were presented. All sessions were recorded using Med Associates Video Monitor software.

#### Stereotaxic Surgery

Viral infusions and optic fiber implant surgeries took place under isoflurane anesthesia (Henry Schein). Mice were anesthetized in an isoflurane induction chamber at 3-4% isoflurane, and then injected with buprenorphine SR (Zoopharm, 0.5 mg/kg s.q.) and carpofen (Zoetis, 5 mg/kg s.q.) prior to the start of surgery. Mice were placed on a stereotaxic frame (Stoetling) and hair was removed from the scalp using Nair. The skin was cleaned with alcohol and a povidone-iodine solution prior to incision. The scalp was opened using a sterile scalpel and holes were drilled in the skull at the appropriate stereotaxic coordinates. Viruses were infused at 100 nl/min through a blunt 33-gauge injection needle using a syringe pump (World Precision Instruments). The needle was left in place for 5 min following the end of the injection, then slowly retracted to avoid leakage up the injection tract. Implants were secured to the skull with Metabond (Parkell) and Flow-it ALC blue light-curing dental epoxy (Pentron). After surgery, mice were allowed to recover until ambulatory on a heated pad, then returned to their homecage with moistened chow or DietGel available. Mice then recovered for three weeks before behavioral experiments began.

#### Fiber Photometry

Mice for fiber photometry experiments received infusions of 1µl of AAV5-CAG-FLEX-jGCaMP7b-WPRE (1.02e13 vg/mL, Addgene, lot 18-429) into lateral SNc (AP -3.1, ML 1.3, DV - 4.2) in one hemisphere and medial SNc (AP -3.1, ML 0.8, DV -4.7) in the other. Hemispheres were counterbalanced between mice. Fiber optic implants (Doric Lenses; 400 μm, 0.48 NA) were placed above DMS (AP 0.8, ML 1.5, DV -2.8) and DLS (AP -0.1, ML 2.8, DV -3.5). The DMS implant was placed in the hemisphere receiving a medial SNc viral injection, while the DLS implant was placed in the hemisphere receiving a lateral SNc viral injection. Calcium signals from dopamine terminals in DMS and DLS were recorded during RI30, on the first and last days of RI60/RR20 training as well as on both footshock probes for each mouse. All recordings were done using a fiber photometry rig with optical components from Doric lenses controlled by a real-time processor from Tucker Davis Technologies (TDT; RZ5P). TDT Synapse software was used for data acquisition. 465nm and 405nm LEDs were modulated at 211 Hz and 330 Hz, respectively, for DMS probes. 465nm and 405nm LEDs were modulated at 450 Hz and 270 Hz, respectively for DLS probes. LED currents were adjusted in order to return a voltage between 150-200mV for each signal, were offset by 5 mA, were demodulated using a 4 Hz lowpass frequency filter. Behavioral timestamps, e.g. for nosepokes and port entries, were fed into the real-time processor as TTL signals from the operant chambers (MED Associates) for alignment with the neural data.

#### Excitatory Optogenetic Stimulation

Mice for DMS (Fig. 5) and DLS (Fig. 7) excitatory optogenetics experiments received 1 µl of AAV5-EF1α-DIO-hChR2(H134R)-EYFP (3.3e13 GC/mL, Addgene, lot v17652) or the control fluorophore-only virus AAV5-EF1α-DIO-EYFP (3.5e12 virus molecules/mL, UNC Vector Core, lot AV4310K) in medial (AP -3.1, ML 0.8, DV -4.7) or lateral SNc (AP -3.1, ML 1.3, DV -4.2) and a single fiber optic implant (Prizmatix; 250µm core, 0.66 NA) over ipsilateral DMS (AP 0.8, ML 1.5, DV -2.8) or DLS (AP -0.1, ML 2.8, DV -3.5). Hemispheres were counterbalanced between mice. During operant training (beginning with FR1), each rewarded nosepoke was paired with a train of blue light (460nm, 1s, 20 Hz, 15 mW) generated by an LED light source and pulse generator (Prizmatix). A subset of mice (“ChR2 Scrambled”) received the same train of light but paired with random nosepokes on a separate RI60 schedule.

#### Inhibitory Optogenetic Stimulation

Mice for DMS inhibitory optogenetics experiments received 1 µl per side of AAV5-EF1α-DIO-eNpHR3.0-EYFP (1.1e13 GC/mL, Addgene, lot v32533) or the control fluorophore-only virus AAV5-EF1α-DIO-EYFP (3.5e12 virus molecules/mL, UNC Vector Core, lot AV4310K) in bilateral medial SNc (AP -3.1, ML 0.8, DV -4.7) and bilateral fiber optic implants (Prizmatix; 500µm core, 0.66 NA) in DMS (AP 0.8, ML ±1.5, DV -2.8). During operant training (beginning with FR1), each rewarded nosepoke was paired with a continuous pulse of orange/red light (625nm, 1s, 15 mW) generated by an LED light source and pulse generator (Prizmatix). A subset of mice (“NpHR Scrambled”) received the same continuous pulse of light but paired with random nosepokes on a separate RI60 schedule.

#### Transcardial Perfusions

Mice received lethal i.p. injections of Euthasol (Virbac, 1mg/kg) a combination of sodium pentobarbital (390 mg/ml) and sodium phenytoin (50 mg/ml), to induce a smooth and rapid onset of unconsciousness and death. Once unresponsive to a firm toe pinch, an incision was made up the middle of the body cavity. An injection needle was inserted into the left ventricle of the heart, the right atrium was punctured and solution (PBS followed by 4% PFA) was infused as the mouse was exsanguinated. The mouse was then decapitated and its brain was removed and fixed overnight at 4°C in 4% PFA.

#### Histology

After perfusion and fixation, brains were transferred to a solution of 30% sucrose in PBS, where they were stored for at least two overnights at 4°C before sectioning. Tissue was sectioned on a freezing microtome (Leica) at 30 μm, stored in cryoprotectant (30% sucrose, 30% ethylene glycol, 1% polyvinyl pyrrolidone in PB) at 4°C until immunostaining. Tyrosine hydroxlase (TH) staining was performed on free floating sections, which were blocked with 3% normal goat serum in PBS-T for 1 hour at room temperature, then stained with 1:500 primary antibody (Aves Labs, Cat No. TYH) in blocking solution at 4°C overnight. Secondary staining was performed using 1:500 goat anti-chicken Alexa Fluor 647 secondary antibody (Life Technologies, Cat. No. A-21449). Anti-GFP staining was performed on free floating sections to amplify signals from GCaMP7b. This staining was performed by blocking in 3% normal goat serum in PBS-T for 1 hour at room temperature, then using 1:500 primary antibody conjugated directly to Alexa Fluor 488 (Life Technologies, Cat. No. A-21311) in blocking solution at 4°C overnight. Tissue was mounted on slides in PBS and coverslips were secured with Fluoromont-G (Southern Biotech). Slides were imaged using a fluorescent microscope (Keyence BZ-X800) with 5x and 40x air immersion objectives. Probe placements were determined by comparing to the Mouse Brain Atlas (Franklin & Paxinos, 2008). GCaMP neurons expressing YFP were counted and colocalized with TH+ neurons using ImageJ software.

### Quantification and Statistical Analysis

#### Behavioral Analysis

Cue-evoked freezing during fear conditioning was scored manually by two blind observers from a recording of the fear conditioning test session using EthoVision software (Noldus). Scores from the two observers were averaged. Freezing was measured throughout the session as a mouse remaining still for more than two seconds. For all other studies, behavioral data was collected automatically by MED-PC software (Med Associates). To sort mice into PR, DPR, and PS groups, we calculated the percent change in shocks received from the early to late shock probe for each mouse. Mice in the top quartile of changers (who increased the number of shocks received by greater than 85%) were classified as delayed punishment resistant (DPR; n=9). The remaining mice were sorted by a median split, with mice receiving more than 13 shocks on the first probe classified as punishment resistant (PR; n=9) and those earning fewer as punishment sensitive (PS; n=18, total n=36, Fig. 2A). The subset of mice that received the omission probe were also sorted by a median split of omission completion time (time to 50 rewards), with mice taking more than 29 minutes classified as long omission (n=10) and those taking less time as short omission (n=10, Fig. S4H).

Plots in Fig. 2O, 5H, 6H, and 7H were generated by plotting a segmental linear regression with lines for the average slope of nosepokes/minute across FR1, RI30, and RI60 training to reveal escalation of nosepoke behavior. The shaded area shows the 95% confidence band surrounding each slope. This analysis was done using GraphPad (Prism) software. Inter-reward intervals were calculated as the time from a rewarded nosepoke to the subsequent rewarded nosepoke (Fig. 2L) on each day of RI60 training. A frequency distribution was created and plotted using GraphPad (Prism) software. Plots in Fig S1B-C were generated by binning the number of nosepokes per five minutes during probe sessions and dividing by nosepokes in the same five-minute bin on the most recent day of RI60/RR20 training.

#### Fiber Photometry Analysis

All analysis was done using custom MATLAB (Mathworks) and Python code. Raw data from 465nm and 405nm channels were passed through a zero-phase digital filter (filtfilt function in Matlab) and a least-squares linear fit (parameters derived with polyfit function) was applied to the 405nm control signal to align it to the 465nm signal. ΔF/F was calculated with the following formula: (465nm signal - fitted 405nm signal) / (fitted 405nm signal). To facilitate comparisons across animals, z-scores were calculated by subtracting the mean ΔF/F calculated across the entire session and dividing by the standard deviation. Peri-stimulus time histograms (PSTHs) were created using the TTL timestamps corresponding to behavioral events. Maximum and minimum peak values and locations from PSTHs in main figures were generated using max and min functions in MATLAB for the 1.5 seconds following behavioral event (ie nosepoke, port entry). AUC was calculated using trap function in MATLAB. We used a customized logic for peak detection in Supplemental Figures 3 and 4 adapted from Holly et al (2019) and Muir et al (2018) (Holly et al., 2019; Muir et al., 2018). Events having amplitudes greater than the summation of a median of 30 seconds moving window and two times median absolute deviation (MADs), were filtered out and the median of the resultant trace was calculated. Peaks having local maxima greater than three times MADs of the resultant trace above the median were considered as events.

#### Statistical Methods

Statistical analysis was done using Prism 9 software (GraphPad). One and two-way ANOVAs, or mixed effects analyses were performed with Tukey’s multiple comparisons and Bonferroni post-hoc analyses when statistically significant main effects or interactions were found. A Kolmogorov-Smirnov test was used to compare the distributions of inter-reward intervals. One RI60 mouse was excluded from the fiber photometry study due to improper fiber placement. A total of six mice were excluded from the optogenetics studies—five due to improper probe placement and one because of illness. All n values listed above do not include these mice.

## Data and Code Availability

The code generated during this study is available at [will make public Github repository on publication].

## SUPPLEMENTAL INFORMATION

Supplemental Information includes seven figures.

---

**Figure S1.**
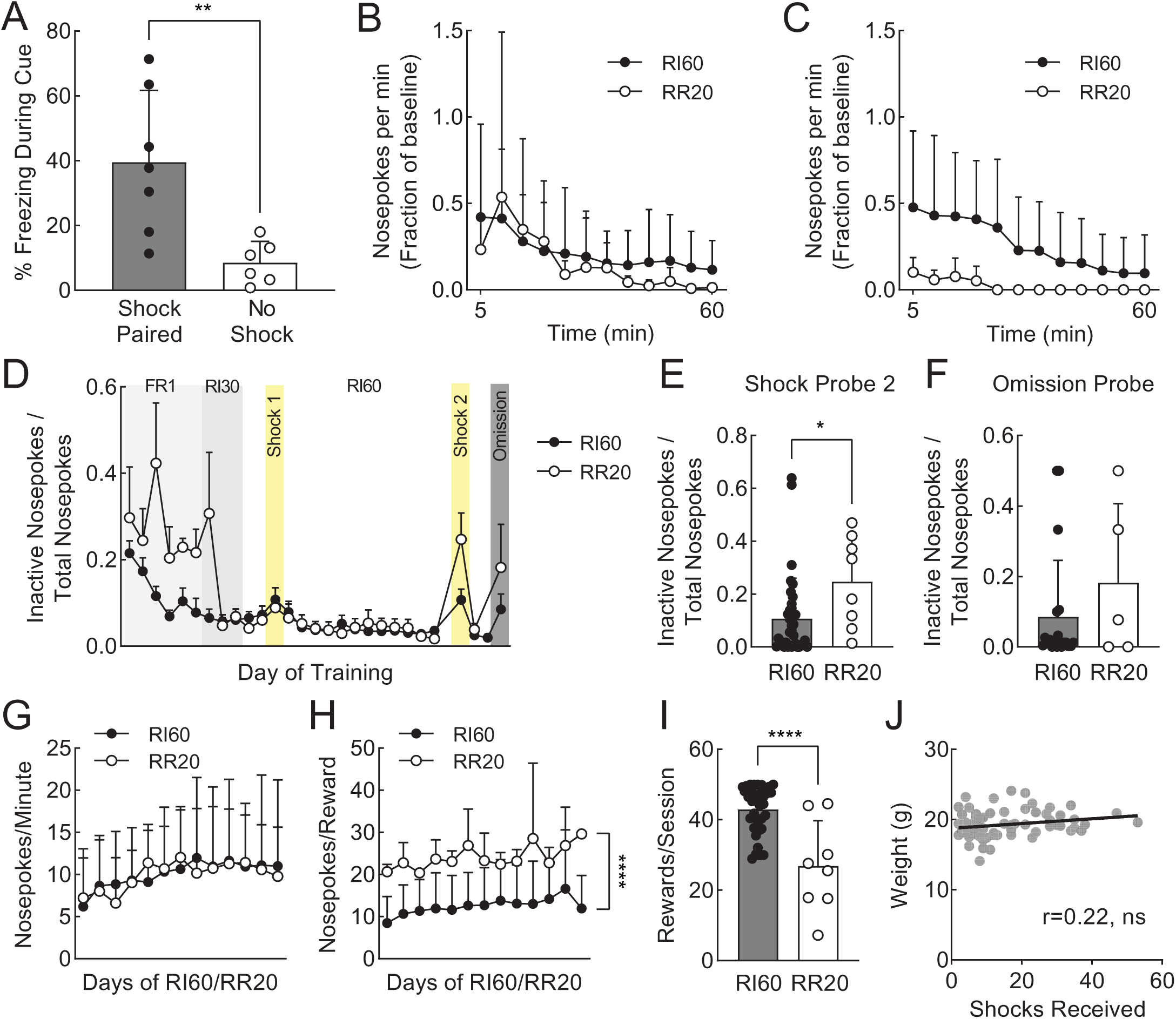
Related to Figure 1. **A.** Percent time spent freezing during a tone cue that had been paired with a 1s 0.2mA shock (black) or no shock (white). Error bars represent SD **p<0.01 **B.** Average nosepokes per minute on the second shock probe represented as a fraction of nosepokes made per minute during the most recent RI60 (black) or RR20 (white) session. Points represent 5 minute bins. Error bars represent SD. **C.** Average nosepokes per min on the omission probe represented as a fraction of nosepokes made per minute during the most recent RI60/RR20 session. Points represent 5 minute bins. Error bars represent SD. **D.** Average nosepokes on the inactive port as a fraction of the total nosepokes across days of training and probe sessions. Error bars represent SD. **E.** Average nosepokes on the inactive port as a fraction of the total nosepokes during the second shock probe session. Error bars represent SD *p<0.05. **F.** Average nosepokes on the inactive port as a fraction of the total nosepokes during the omission probe session. Error bars represent SD. **G.** Average nosepokes per minute across days of RI60/RR20 training. Error bars represent SD. **H.** Average nosepokes per reward earned across days of RI60/RR20 training. Error bars represent SD. **I.** Average rewards earned per session of RI60/RR20 training. Error bars represent SD ****p<0.0001. **J.** Correlation between weight of animals (grams) and number of shocks received during shock sessions for RI60-trained animals (r=0.22, ns).

**Figure S2.**
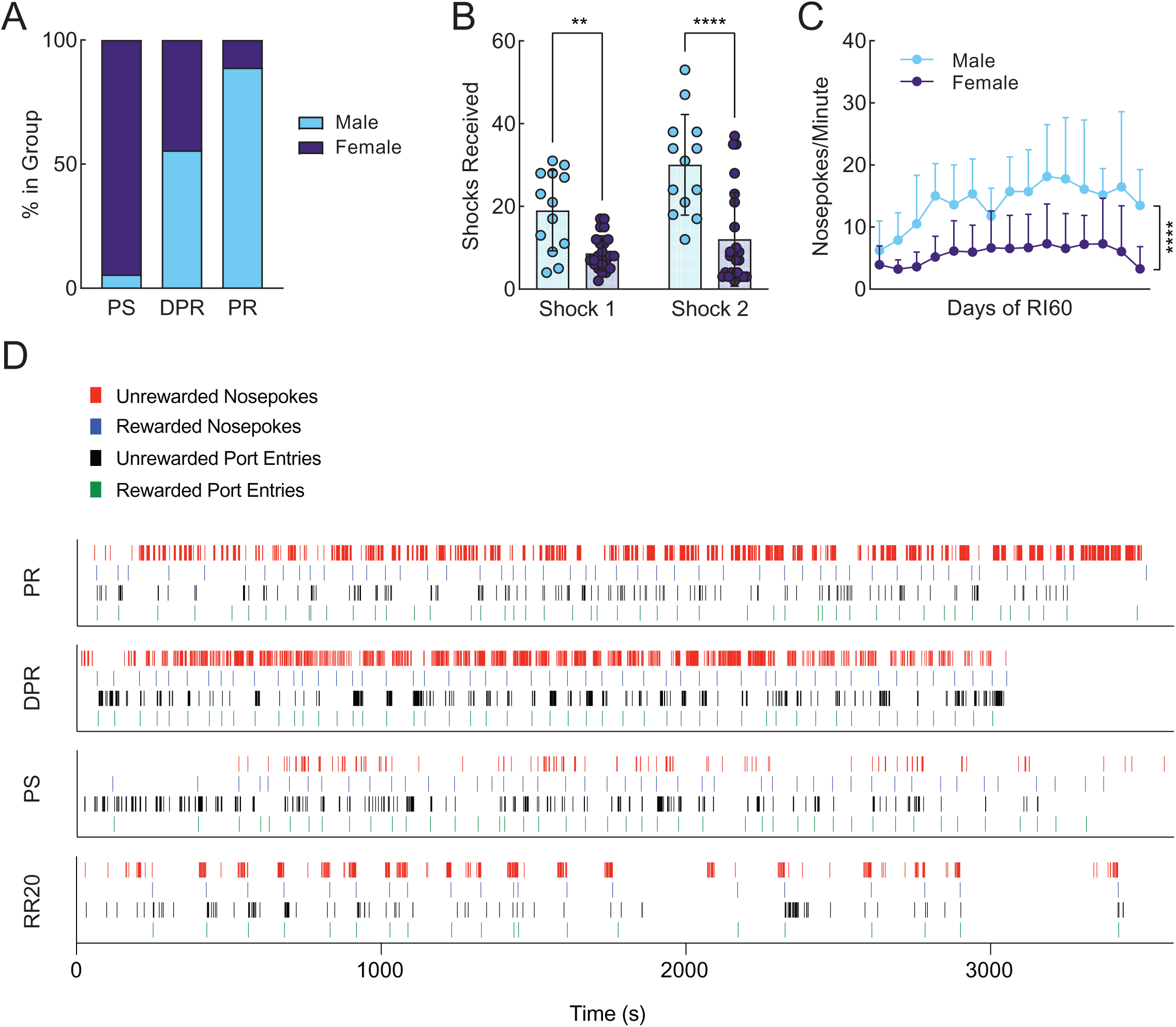
Related to Figure 2. **A.** Percentage of male (blue) and female (purple) mice in each behavioral phenotype. **B.** Average shocks received on early and late shock probes for each sex. Error bars represent SD **C.** Average nosepokes per minute across days of RI60 training. Error bars represent SD ****p<0.0001. **D.** Examples of behavioral timestamps recorded over the course of a full RI60 or RR20 session for a representative mouse from each group.

**Figure S3.**
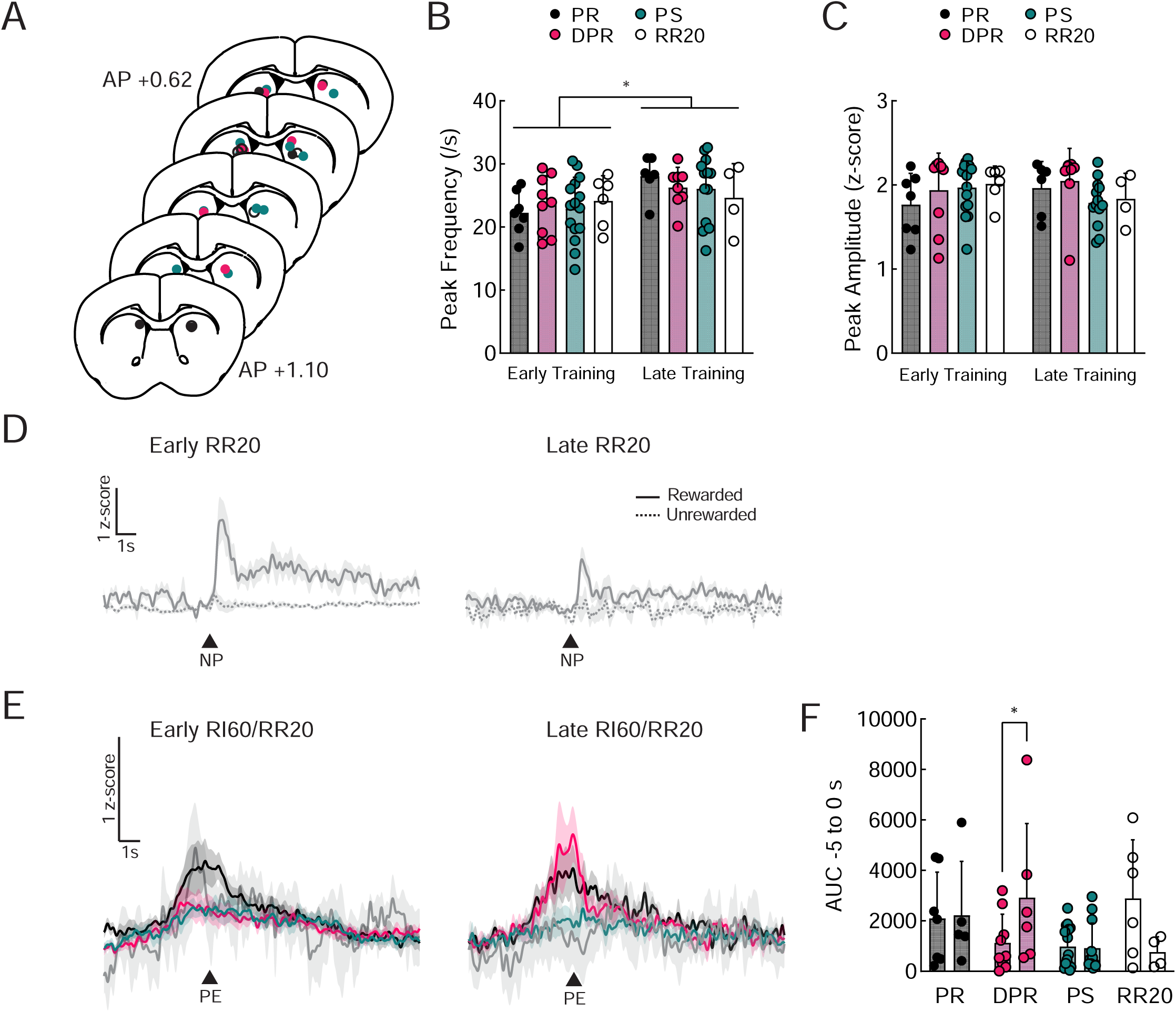
Related to Figure 3. **A.** Probe placements in DMS for all mice included in Figure 3. **B.** Average frequency of peaks detected in the DMS dopamine terminal signal for all groups. Error bars represent SD *p<0.05. Punishment resistant (PR; black), delayed punishment resistant (DPR; pink), or punishment sensitive (PS; teal), RR20-trained (RR20; white). **C.** Average amplitude of detected peaks in the DMS dopamine terminal signal for all groups. Error bars represent SD. **D.** Peri-stimulus time histograms (PSTHs) showing the average signal from DMS dopamine terminals at the time of rewarded (solid) and unrewarded (dashed) nosepokes (NP) for RR20-trained mice early in training (RR20 day one or two) and late in training (RR20 day eleven or twelve). Shaded region represents SEM. **E.** Peri-stimulus time histograms (PSTHs) showing the average signal from DMS dopamine terminals at the time of rewarded port entry (PE) for each phenotype early in training (RI60 day one or two) and late in training (RI60 day eleven or twelve). Shaded region represents SEM. **F.** Quantification of average area under the curve (AUC) of the DMS dopamine terminal signal from -5 to 0 seconds, relative to a rewarded port entry. Error bars represent SD *p<0.05.

**Figure S4.**
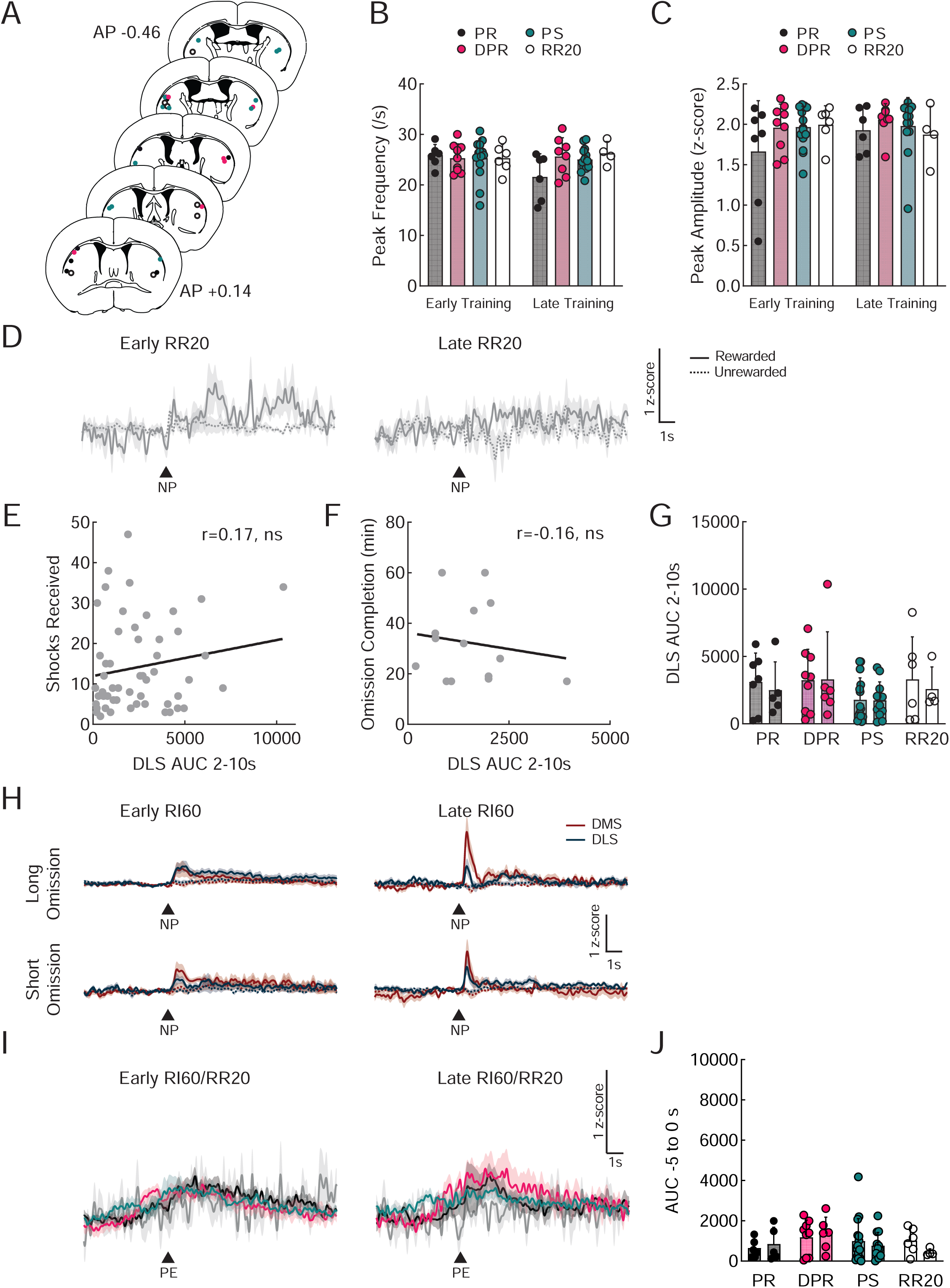
Related to Figure 4. **A.** Probe placements in DLS for all mice included in Figure 4. **B.** Average frequency of peaks detected in the DLS dopamine terminal signal for all groups. Error bars represent SD. Punishment resistant (PR; black), delayed punishment resistant (DPR; pink), or punishment sensitive (PS; teal), RR20-trained (RR20; white). **C.** Average amplitude of detected peaks in the DLS dopamine terminal signal for all groups. Error bars represent SD. **D.** Peri-stimulus time histograms (PSTHs) showing the average signal from DLS dopamine terminals at the time of rewarded (solid) and unrewarded (dashed) nosepokes (NP) for RR20-trained mice early in training (RR20 day one or two) and late in training (RR20 day eleven or twelve). Shaded region represents SEM. **E.** Correlation of shocks received and average area under the curve of DLS dopamine terminal signal from 2 to 10s, relative to the rewarded nosepoke (r=0.17, ns). **F.** Correlation of omission completion time and average area under the curve of DLS dopamine terminal signal from 2 to 10s, relative to the rewarded nosepoke (r=-0.16, ns). **G.** Quantification of average area under the curve (AUC) of the DLS dopamine terminal signal from 2 to 10 seconds, relative to the rewarded nosepoke. Error bars represent SD. **H.** Peri-stimulus time histograms (PSTHs) showing the average signal from DMS (red) and DLS (blue) dopamine terminals at the time of rewarded (solid) and unrewarded (dashed) nosepokes (NP) for animals sorted by omission completion time early in training (RI60 day one or two) and late in training (RI60 day eleven or twelve). Shaded region represents SEM. **I.** Peri-stimulus time histograms (PSTHs) showing the average signal from DLS dopamine terminals at the time of rewarded port entry (PE) for each phenotype early in training (RI60 day one or two) and late in training (RI60 day eleven or twelve). Shaded region represents SEM. **F.** Quantification of average area under the curve (AUC) of the DLS dopamine terminal signal from -5 to 0 seconds relative to a rewarded port entry. Error bars represent SD.

**Figure S5.**
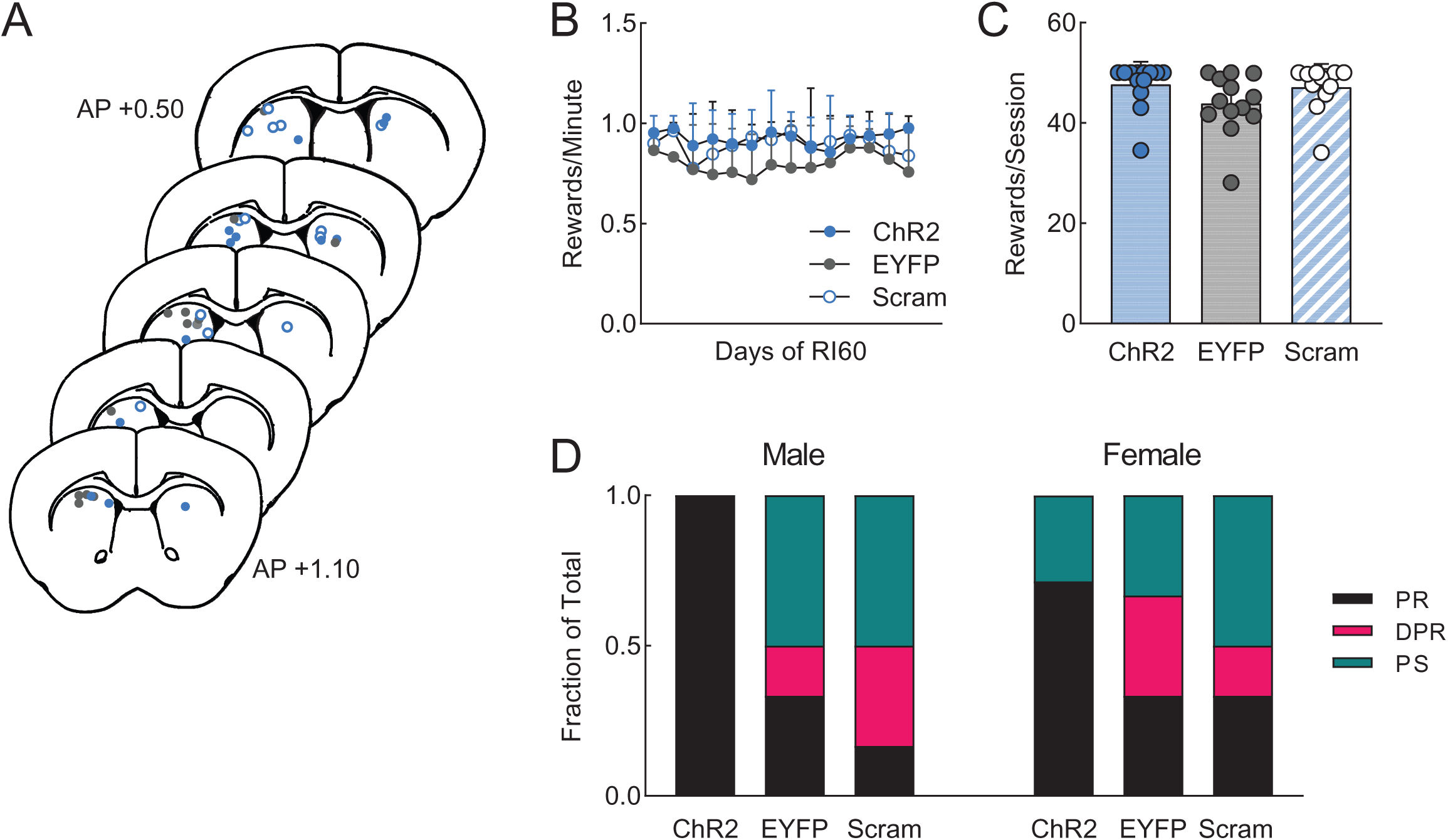
Related to Figure 5. **A.** Probe placements in DMS for all mice included in Figure 5. ChR2 (blue), EYFP (gray), ChR2 scrambled (blue outline). **B.** Average number of rewards earned per minute across days of RI60 training. Error bars represent SD. **C.** Average rewards earned per day of RI60 training. Error bars represent SD. **D.** Fraction of each behavioral phenotype (punishment resistant=black, delayed punishment resistant=pink, punishment sensitive=teal) in each manipulation divided by sex.

**Figure S6.**
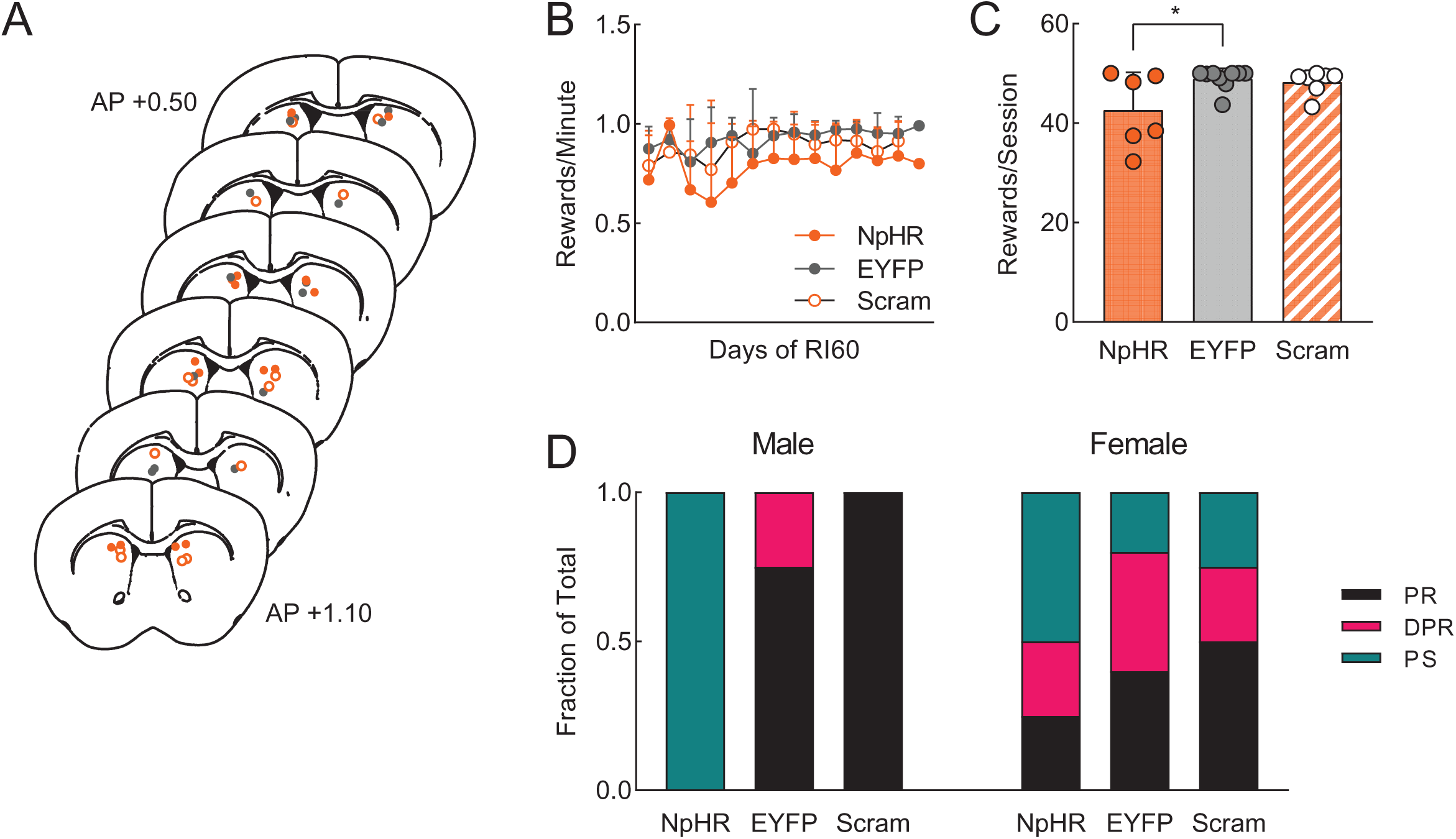
Related to Figure 6. **A.** Probe placements in DMS for all mice included in Figure 6. ChR2 (orange), EYFP (gray), ChR2 scrambled (orange outline). **B.** Average number of rewards earned per minute across days of RI60 training. Error bars represent SD. **C.** Average rewards earned per day of RI60 training. Error bars represent SD *p<0.05. **D.** Fraction of each behavioral phenotype (punishment resistant=black, delayed punishment resistant=pink, punishment sensitive=teal) in each manipulation divided by sex.

**Figure S7.**
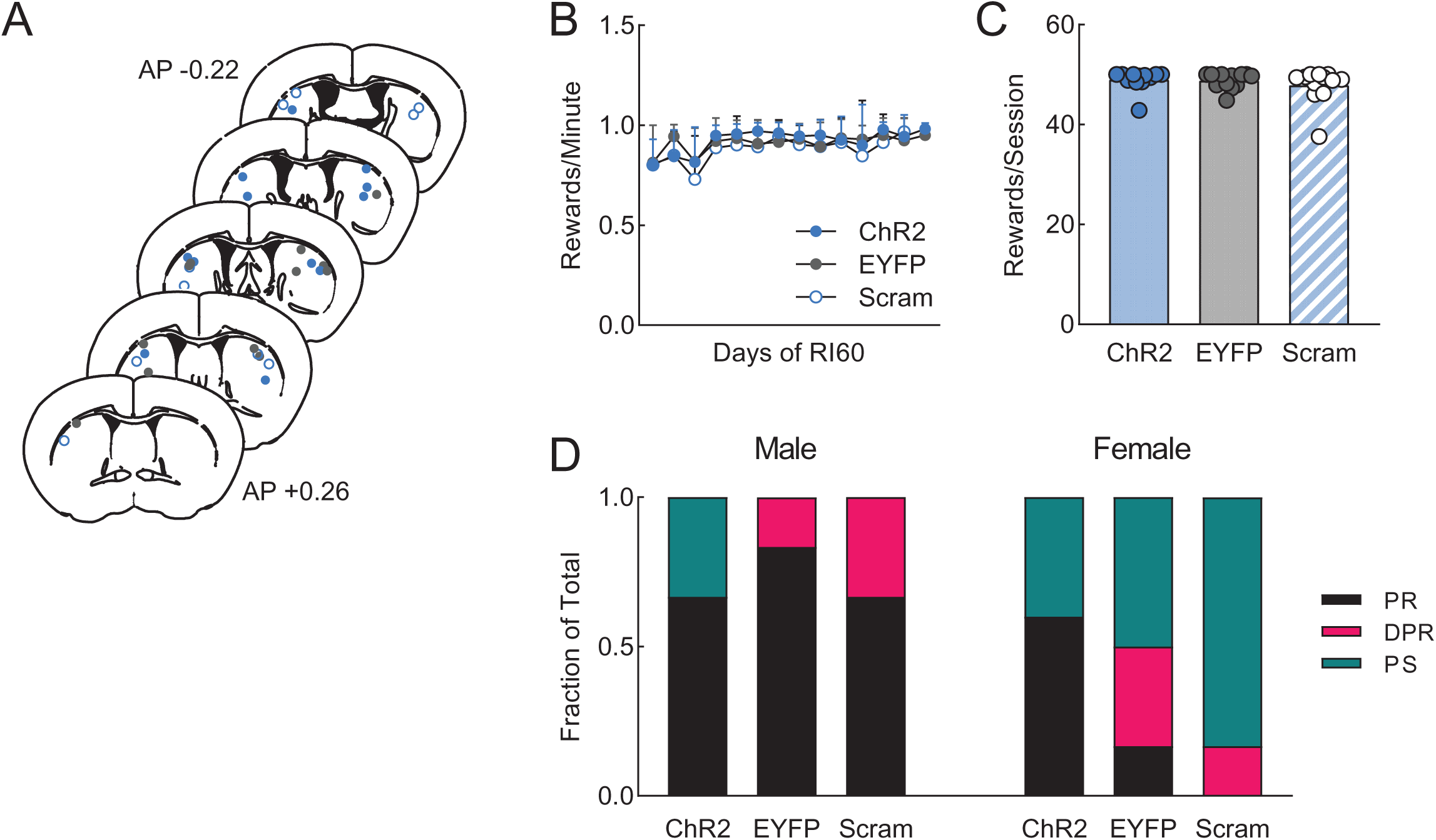
Related to Figure 7. **A.** Probe placements in DLS for all mice included in Figure 7. ChR2 (blue), EYFP (gray), ChR2 scrambled (blue outline). **B.** Average number of rewards earned per minute across days of RI60 training. Error bars represent SD. **C.** Average rewards earned per day of RI60 training. Error bars represent SD. **D.** Fraction of each behavioral phenotype (punishment resistant=black, delayed punishment resistant=pink, punishment sensitive=teal) in each manipulation divided by sex.

